# An essential role for MEF2C in the cortical response to loss of sleep

**DOI:** 10.1101/2020.04.27.064535

**Authors:** Theresa E. Bjorness, Ashwinikumar Kulkarni, Volodymer Rybalchenko, Ayako Suzuki, Catherine Bridges, Adam J. Harrington, Christopher W. Cowan, Joseph S. Takahashi, Genevieve Konopka, Robert W. Greene

## Abstract

Neuronal activity and gene expression in response to the loss of sleep can provide a window into the enigma of sleep function. Sleep loss is associated with brain differential gene expression, an increase in pyramidal cell mEPSC frequency and amplitude, and a characteristic rebound and resolution of slow wave sleep-slow wave activity (SWS-SWA). However, the molecular mechanism(s) mediating the sleep loss response are not well understood. We show that sleep-loss regulates MEF2C phosphorylation, a key mechanism regulating MEF2C transcriptional activity, and that MEF2C function in postnatal excitatory forebrain neurons is required for the biological events in response to sleep loss. These include altered gene expression, the increase and recovery of synaptic strength, and the rebound and resolution of SWS-SWA, which implicate MEF2C as an essential regulator of sleep function.

**One Sentence Summary:** MEF2C is critical to the response to sleep loss.

## Main Text

Sleep abnormalities are commonly observed in numerous neurological disorders, including autism spectrum disorder, major depressive disorder, bipolar disorder, post-traumatic stress disorder, neurodegenerative disorders and many others, but our understanding of sleep need and its regulation and resolution is poorly understood. Following an extended period of waking, or sleep deprivation (SD), the mammalian cortex shows an altered pattern of EEG activity characterized by rebound slow wave power (delta power in the frequency range of 0.5-4.5Hz) during the ensuing slow wave sleep (SWS; also referred to as NREM) periods. The amplitude of the rebound directly correlates with the previous waking or SD duration (Borbely, 1982; Franken, Chollet, & Tafti, 2001) implicating rebound SWS, slow wave activity (SWS-SWA) as a biomarker of sleep need. During SWS episodes, the rebound SWS-SWA increase resolves, consistent with resolution of sleep need (Bjorness et al., 2016).

The buildup and resolution of sleep need is correlated with overall cortical, excitatory synaptic strength. Based on sleep-related modulation of excitatory synapse biochemistry, morphology and electrophysiological activity, a scaling-up during waking and a scaling-down during sleep have been observed (Bushey, Hughes, Tononi, & Cirelli, 2010; de Vivo et al., 2017; Diering et al., 2017; Liu, Faraguna, Cirelli, Tononi, & Gao, 2010), although this may not necessarily be reflected by the neuronal firing rates (Hengen, Torrado Pacheco, McGregor, Van Hooser, & Turrigiano, 2016; Watson, Levenstein, Greene, Gelinas, & Buzsaki, 2016).

Changes in synaptic transcript (Noya et al., 2019) and protein (Diering et al., 2017; Noya et al., 2019) expression, required for synaptic down-scaling, are observed during the sleep phase in association with an average decrease in synapse size (de Vivo et al., 2017; Diering et al., 2017). Characterization of the cell-signaling biochemical and molecular mechanisms responsible for these sleep-related changes in protein expression is needed.

In this study, we examined neurobiological changes in the mouse frontal cortex in response to sleep loss, involving transcriptomic architecture, regulation of synaptic strength in excitatory cortical neurons and SWS-SWA, correlated with sleep need. The MEF2 transcription factors (Flavell et al., 2006) and, more particularly, the transcription factor MEF2C (Harrington et al., 2016; Li et al., 2008; Rajkovich et al., 2017), modulate synaptic strength in excitatory cortical neurons in response to neuronal activity, reviewed in (Assali, Harrington, & Cowan, 2019). As changes in neuronal activity may be observed across sleep-wake states, we investigated the role of *Mef2c* in sleep-related control of gene expression, sleep-related modulation of synaptic function and sleep-need correlated SWS-SWA buildup and resolution.

## Results

### Wake-sleep mediated control of the transcriptome

To evaluate the effects of sleep loss, we compared three sleep-related conditions for differentially expressed genes (DEGs) from samples of the frontal cortex (FC, the cortical region showing the highest power of SWS-SWA) of male C57BL/6J (WT) mice (Fig. 1A) including: 1) control sleep (CS; mice free to sleep, ad libitum; n=6), 2) sleep deprived for 6h (SD; n=7), and 3) sleep deprived for 4h followed by recovery sleep for 2h (RS; n=7). All samples were collected 6h after lights on (ZT6), controlling for circadian influence, and were processed for RNA-seq analysis (see Materials and Methods).

**Fig. 1.**
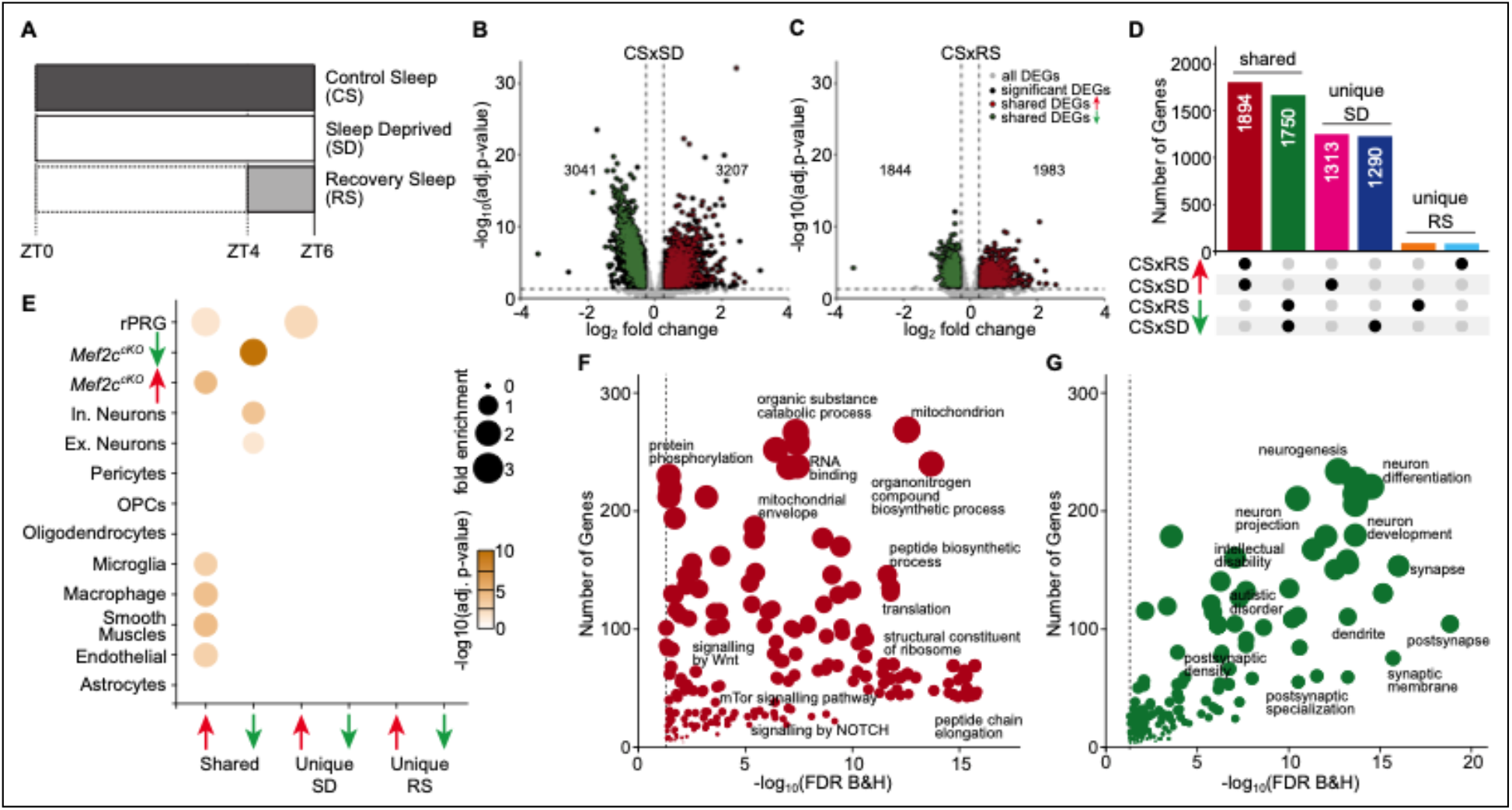
Sleep need induced transcriptomic changes. (**A**) Schematic of the experimental design illustrating the protocol for frontal cortex brain tissue collection from three sleep conditions at Zeitgeber Time (ZT)=6 hours: control sleep (CS), sleep deprived (SD) and recovery sleep (RS). Volcano plots showing differentially expressed genes across (**B**) CS to SD and (**C**) CS to RS. Significantly differentially expressed genes (DEGs) (adj. P-value <= 0.05, absolute log_2_ fold change >= 0.3), unique to the sleep condition, are indicated as *black* dots whereas genes shared by SD and RS are indicated as either *red* (increased expression) or *green* (decreased expression) dots. (**D**) Upset plot showing the shared and unique sets of significant DEGs for CS to SD and CS to RS. (**E**) Dot plot showing enrichment of significant DEGs for cell-types identified using single cell RNA sequencing data (Hrvatin et al., 2018) from mouse cortex, identified *Mef2c* target genes (Harrington et al., 2016) and rapid primary response genes (rPRGs) (Tyssowski et al., 2018). Dot plot showing GO analysis for shared significant DEGs with (**F**) increased and (**G**) decreased expression across SD and RS compared to CS (for complete list of significant GO, see Table S3).

The CS samples expressed, on average, ~13,000 genes with CPM≥1.0 in the FC. Comparing the SD samples to the CS samples, we observed 6,248 DEGs with roughly half the number of genes having decreased expression (FDR<= 0.05; |log_2_ (fold change)|>=0.3; Fig.1B; Table S1), confirming an earlier report of massive transcriptome changes associated with SD (Diessler et al., 2018). Similarly, the RS cohort showed 3827 DEGs compared to CS, again with approximately half the number of genes showing decreased expression (Fig. 1C; Table S1).

In contrast, only 81 genes showed significantly altered expression comparing the SD condition to RS (Table S1), suggesting that for the majority of SD-sensitive genes, RS is more than simply a direct recovery of altered gene expression back to CS levels. Rather, more than half of the differential expression initiated during SD is continued during RS (Fig. 1D), significantly extending the SD-induced change.

Because SD may be considered stressful, we compared classic glucocorticoid-mediated response genes to the increased DEGs observed in response to SD (Fig. S1, Table S2). No significant increase in glucocorticoid (GC) intracellular response genes was found in the up-regulated SD DEGs (p-value=0.513 using Fisher’s exact test), suggesting a more SD-specific response than other GC-mediated stress responses.

Using previously published single cell RNA-seq from the adult mouse cortex (Hrvatin et al., 2018), we identified cell type-specific gene expression patterns across sleep conditions (Fig. 1E). Interestingly, the down regulated DEGs shared by SD and RS were enriched for genes associated with both excitatory and inhibitory neurons, whereas the upregulated DEGs shared by SD and RS were enriched for genes associated with nonneuronal cell types (Fig. 1E).

Since many of the MEF2C-regulated genes expressed in CNS neurons respond to neuronal activation with downregulation of excitatory synaptic strength (Flavell et al., 2006; Rajkovich et al., 2017), we also examined MEF2C target gene expression in response to SD or RS. We found that the downregulated DEGs shared by SD and RS were enriched for neuronal genes downregulated by *Mef2c* loss of function as previously observed in a conditional knockout of neuronal *Mef2c, Mef2c^Emx1-cKO^* (Harrington et al., 2016). Whereas the upregulated DEGs in response to SD and RS were enriched for *Mef2c* non-neuronal genes (Fig. 1E). Thus, the transcriptome changes induced by SD and RS in the cortex mimic those seen with loss of MEF2C.

We next functionally characterized the sleep-loss induced transcriptional changes shared by SD and RS states by carrying out a gene ontology (GO) analysis. Overall, there are distinct GO categories for the upregulated and downregulated genes (Fig. 1F, G; Table S2). The upregulated categories include mitochondrial components and signaling pathways of oxidative phosphorylation and anaplerosis (Fig. S2), indicative of energy mobilization, as well as anabolic protein metabolism that includes ribosomal components together with transcriptional, translational and related processes. These results are consistent with and extend those from a previous study using microarrays (Cirelli, Gutierrez, & Tononi, 2004) that lead to a conclusion, “… that sleep, far from being a quiescent state of global inactivity, may actively favor specific cellular functions”. Recently, similar findings were reported in a RNAseq analysis from C57BL/6J whole cortex and a “gentle handling” method of SD (Hor et al., 2019).

Several studies provide converging evidence for a general decrease in cortical excitatory synaptic strength in association with sleep (de Vivo et al., 2017; Liu et al., 2010; Maret, Faraguna, Nelson, Cirelli, & Tononi, 2011) with respect to protein expression in synaptosomes (Diering et al., 2017) and spontaneous miniature excitatory post-synaptic currents and frequency (Liu et al., 2010; Vyazovskiy, Cirelli, Pfister-Genskow, Faraguna, & Tononi, 2008). Consistent with this, we observed that the downregulated genes observed in SD and RS are enriched for GO categories reflecting synaptic biological processes and cellular components. These genes are also enriched for genes involved in diseases of cognition associated with abnormal synaptic function, including schizophrenia, autism, intellectual disability and neurodegenerative disorders (Fig. 1G, Table S3).

Some of the SD-induced DEGs have a notable role as both transcription factors and/or first responders to neural activity and sensory stimulation. Rapid, primary response genes (rPRG) have been identified in cortical neuronal cultures as responders to both brief or 6 h exposures to a depolarizing concentration of potassium in the presence of a translational blocker (Tyssowski et al., 2018). We find that DEGs for CS compared to SD and/or RS conditions are significantly enriched for these rPRGs (Fig. 1E). Furthermore, early response transcription factors (ERTF;(Hrvatin et al., 2018)) were identified in visual cortex due to their response to a short exposure to light after being in constant darkness. We observed that 50% of these were DEGs for CS to SD (listed in Table S1). Notably, these ERTFs from the FC responded to 6h SD, carried out in constant darkness, suggesting that SD alone is a sufficient stimulus for their activation and raising the possibility that their targets in response to SD are different from those identified by brief light exposure (Fowler, Sen, & Roy, 2011).

It was recently shown that the influence of circadian rhythms on gene transcript expression in synaptosomes was relatively unaffected by SD, although SD did prevent translation (Noya et al., 2019). Nevertheless, many core clock genes are known to affect sleep homeostasis and the expression of some core clock genes, in particular, *Per2*, are altered by SD, as previously reviewed (Franken, 2013). To control for circadian-mediated changes in gene expression, we assessed DEGs at the same circadian time, ZT6, changing only the 6 h, immediate sleep history. In contrast to DEGs in synaptosomes, we observed in FC tissue that SD from ZT=0 to 6h resulted in a number of core clock DEGs (compared to CS; Fig. S3), including downregulation of *Arntl, Clock* and *Npas2* and downregulation of a set of genes regulating circadian entrainment (MGI; GO:0009649). Taken together, these data are consistent with an SD-induced attenuation and destabilization of circadian-mediated gene expression and confirm recent observations using an RNAseq assessment in time series together with SD (Hor et al., 2019).

To further uncover the organizational architecture and prioritize the extensive SD-mediated transcriptome, we used weighted gene co-expression network analysis (WGCNA), with expressed genes from all three conditions (CS, SD, and RS) to generate a network of 59 modules. Many of the modules (53%) were enriched for either up or downregulated DEGs from CS to SD conditions (Fig. 2A). Most of those enriched for upregulation from CS to SD (n=11), were also enriched for upregulation from CS to RS (n=10), and similarly, most modules enriched for downregulation from CS to SD (n=18) were also enriched for downregulation from CS to RS (n=13), showing an organization primarily driven by DEGs, that is consistent between SD and RS conditions. The GO of these shared modules (Table S4) is also consistent with those seen using only the lists of DEGs.

**Fig. 2.**
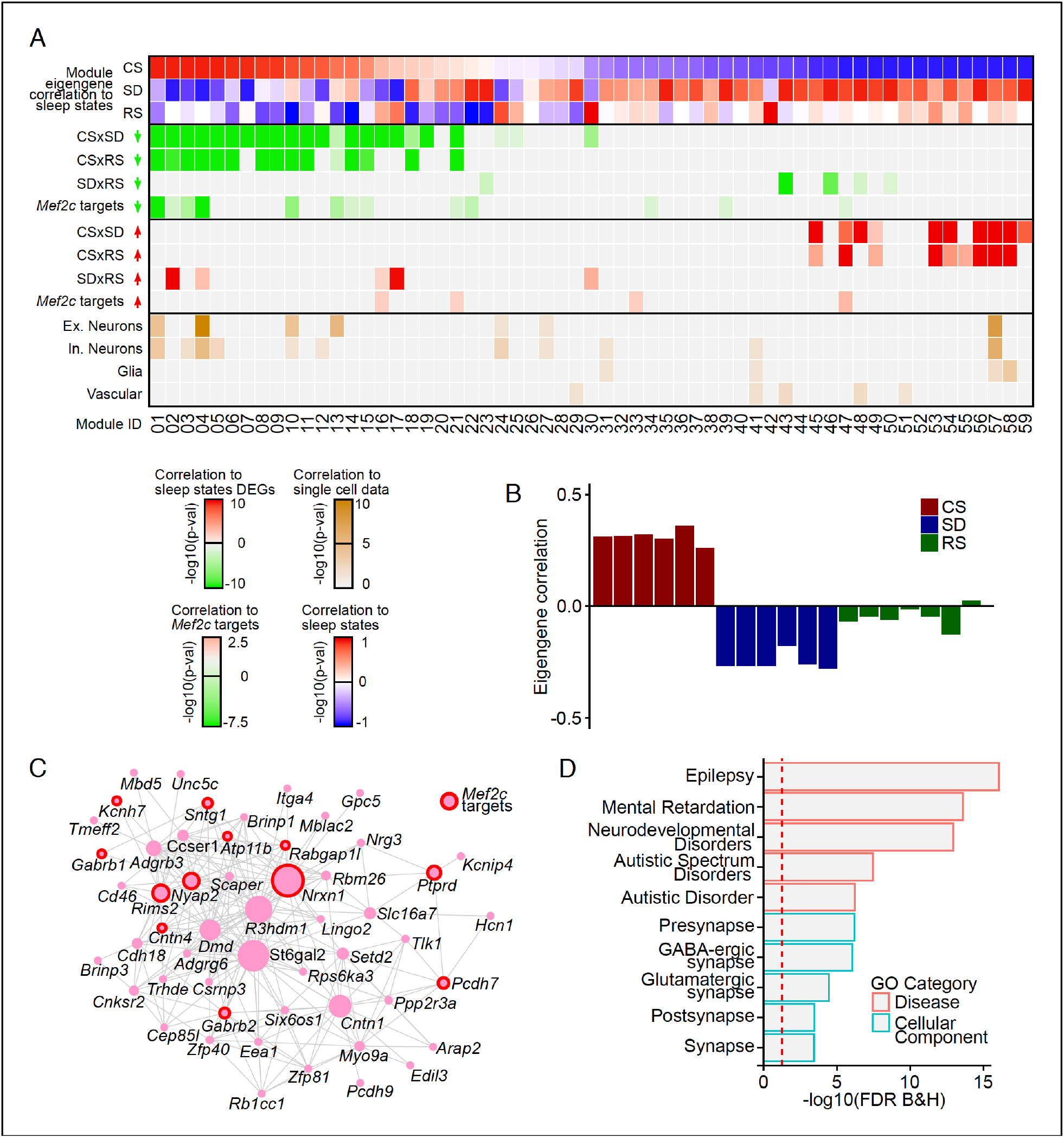
Coordinated transcriptomic responses with sleep need indicate a role for MEF2C. (A) Heatmap showing enrichment of sleep-state DEGs across modules of genes identified by weighted gene co-expression network analysis (WGCNA). First three rows show module eigengene correlation to sleep states (CS, SD and RS respectively) ordered by most positive to most negative correlation to CS sleep state. The following four rows highlight the significance of module enrichment for a specific set of DEGs colored by *green* for genes with decreased expression for CSxSD, CSxRS, SDxRS and MEF2C targets (Harrington et al., 2016) and the next four rows colored by *red* to highlights the significance of module enrichment for genes with increased expression for CSxSD, CSxRS, SDxRS and MEF2C targets (Harrington et al., 2016). The bottom four rows colored by shades of *orangeyellow* show enrichment of modules for cell types identified using single cell RNA sequencing (Hrvatin et al., 2018) from mouse cortex. Each column is a module of genes as indicated by the module ID in last row. (B) Barplot for a selected module (Module ID 04) with eigengene correlation across sleep states showing a strong positive correlation to CS and negative correlation to SD and weak negative correlation to RS. (C) Network plot for selected module (Module ID 04) showing the top 250 connections between genes. Genes that are MEF2C targets (Harrington et al., 2016) are highlighted by red circles. (D) GO plot for genes within the selected module (Module ID 04).

The advantage of WGCNA is that it allows for the prioritization of genes that may be central drivers of gene expression independent of arbitrary cutoffs of p-values and fold changes. We identified two modules (M04 and M10) whose module eigengenes were negatively correlated with SD and RS samples (Fig. 2B and Fig. S4). These two modules were of interest due to enrichment of several major categories of genes (Fig. 2A): 1) downregulated MEF2C target genes, 2) excitatory and inhibitory neuronal genes, 3) genes downregulated in SD and RS, and 4) genes involved in synaptic function, epilepsy and cognitive pathologies including autism (Fig. 2D). These characteristics are consistent with modulation by the transcription factor encoded by MEF2C. Both of these modules have hub genes encoding receptors, ionic channels or cell adhesion, and identified as autism and epilepsy risk factors in addition to being target genes for MEF2C (for complete GO of M04 and M10 and other modules, see Table S4).

#### The role of MEF2C in the transcriptomic response to sleep loss

DEGs comparing CS to SD conditions are enriched for MEF2C targets that down regulate genes controlling synaptic strength in neurons (Fig. 1E, G, Table S3). Modules of correlated gene expression across sleep conditions showed a similar enrichment of MEF2C targets, pattern of expression and GO as the DEGs across sleep conditions (Fig. 2A, Table S5).

To better understand the role of MEF2C across sleep conditions, we examined a conditional knockout mouse using a Cre-loxP system. We crossed a *Mef2c^f/f^* mouse line (Arnold et al., 2007; Zang et al., 2013) with a transgenic *Cre:CamKII* line and examined *Mef2c^cKO^* mice (see methods for details) for their response to sleep loss. *Mef2c^f/f^* mice showed 767 DEGs within 12,403 total expressed genes from CS (n=4) to SD (n=3); whereas, strikingly, the *Mef2c^cKO^* showed only 36 DEGs within 12,620 total expressed genes in CS (n=3) to SD (n=3; Fig. 3A,B,C). Both the *Mef2c^f/f^* and the *Mef2c^cKO^* genotypes expressed similar numbers of FC genes as the WT, using a CPM ≥1.0, but the responses of Mef2^cKO^ to sleep conditions were attenuated to ~7% of the *Mef2c^f/f^* responses, indicating the essential role of *Mef2c* in the transcriptome response to sleep loss.

**Fig 3.**
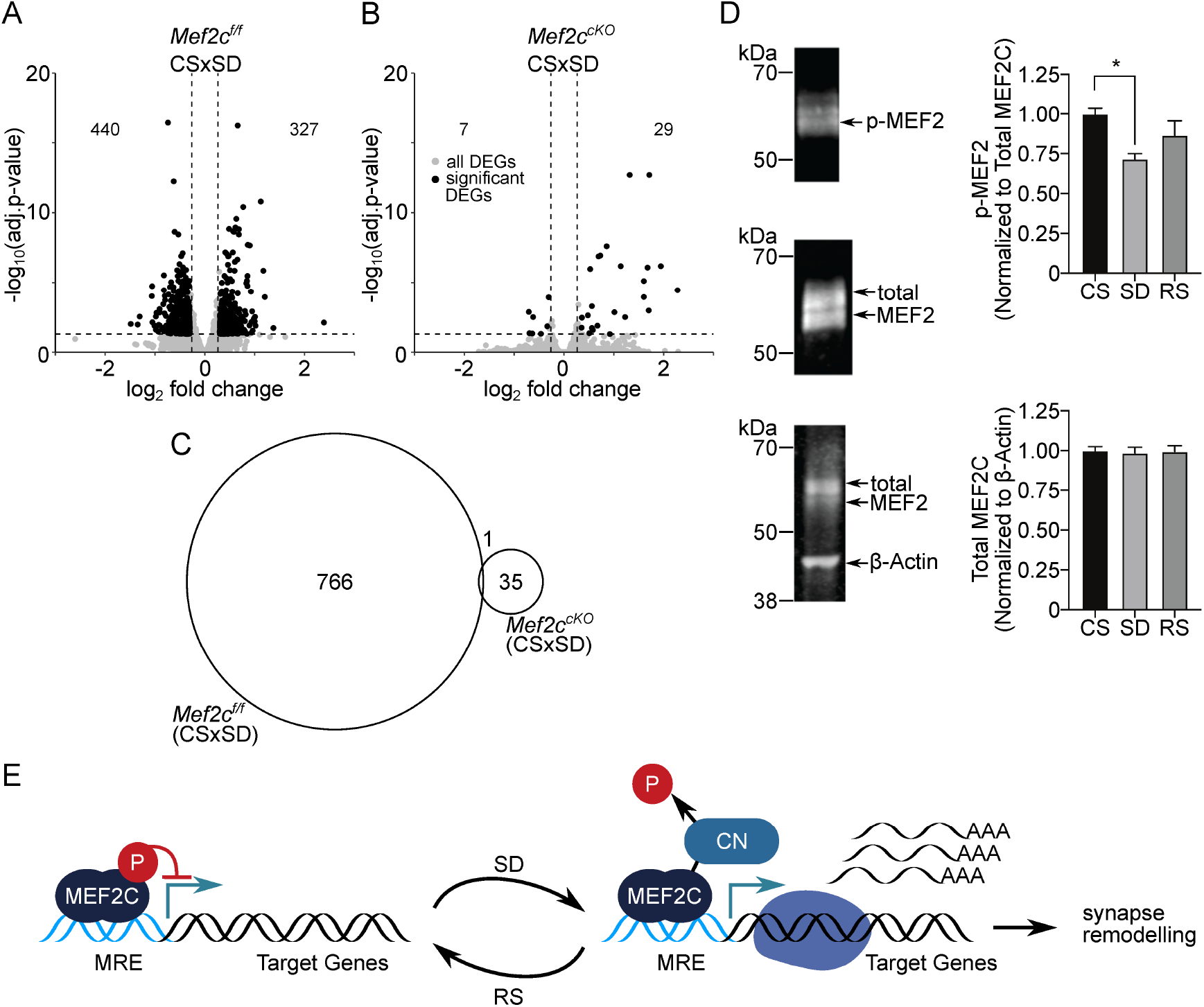
MEF2C, its de-phosphorylation and essential role in sleep loss mediated differential gene expression. (**A**) Volcano plot showing the DEGs identified for Mef2c^f/f^ and another, (**B**) for Mef2c^cKO^ between CS to SD sleep condition. DEGs (adj. P-val <=0.05; absolute log_2_(change)=>0.3) are indicated by black dots. (**C**) A Venn diagram showing overlap between DEGs for Mef2c^f/f^ and Mef2c^cKO^ between CS and SD sleep states. (**D**) Immunoprecipitation of MEF2C to detect phospho-MEF2 and total MEF2C for each sleep condition. Phospho-MEF2 (top blot) re-labelled from immunoprecipitation of total MEF2C (middle blot) was quantified and normalized to total MEF2C signal for each sample (n=4 samples/condition). Total MEF2C from total cell lysate was quantified and normalized to B-actin (bottom blot). Data reported as mean +/- SEM. Statistical significance was determined by one-way ANOVA with Tukey’s multiple comparisons test (interaction = p<0.05, post hoc test comparing CS to SD * = p<0.05). (**E**) Model showing role of de-phosphorylated, activated MEF2C, increased relative to total MEF2C by loss of sleep and leading to synapse remodeling. MRE = Mef2 response element, CN = calcineurin.

Because MEF2 activity is regulated at least in part by its phosphorylation state (Flavell et al., 2006) then it might be expected to change in response to sleep loss and recovery sleep. We examined the phosphorylation state of MEF2C in FC in three sleep conditions (*i.e*. CS, SD and RS) and found that compared to CS, SD produced a significant reduction of phospho-MEF2C (Fig. 3D), but without altering total MEF2C protein levels (Fig. 3D). In contrast, RS did not significantly alter phospho-MEF2C levels (Fig. 3D), suggesting that SD triggers signaling events to transiently activate MEF2C-dependent transcription and promote the weakening and/or elimination of excitatory synapses.

As noted above (Fig. 1E), CS to SD and RS downregulated DEGs were enriched for genes that were downregulated in an earlier study comparing WT to *Mef2c^cKO^* in control conditions (Harrington et al., 2016). Similarly, the upregulated genes from CS to SD and RS are enriched for up-regulated WT to *Mef2c^cKO^* genes. In other words, the DEGs of *Mef2c^cKO^* (compared to WT) in control conditions are similar to the DEGs of CS to SD in WT mice. However, in the *Mef2c^cKO^* used in our study, there is little response to SD (Fig. 3 B,C), rather the DEGs of the WT sleep loss response are now differentially expressed in CS conditions in *Mef2c^cKO^* mice, potentially, a compensatory response, no longer responsive to sleep loss.

#### The role of *Mef2c* in the synaptic response to sleep loss

Based on our transcriptome analyses, SD and RS induced many DEGs affecting synaptic strength and synaptic cellular components that required the MEF2C transcription factor (Fig. 1, 2, 3). MEF2C can regulate forebrain excitatory synapse strength of pyramidal neurons by decreasing synaptic density (Harrington et al., 2016) and frequency (Flavell et al., 2006). Down-scaling of cortical synaptic strength is associated with sleep (de Vivo et al., 2017; Diering et al., 2017). Previous observations indicate that acute loss of sleep is associated with increased miniature excitatory post-synaptic (mEPSC) frequency and amplitude and that recovery sleep reverses these changes in cortical neurons recorded from C57BL/6 mouse FC (Liu et al., 2010). However, neither the involvement of changes in probability of release nor changes in active excitatory synapse number and the role of *Mef2c* have been previously considered.

To directly examine sleep-associated changes in synaptic strength, the same conditions as employed for the transcription analyses (i.e., CS, SD, RS) were applied to *Mef2c^f/f^* male mice. Acute *ex-vivo* slices of FC were prepared from these mice at ZT=6 hours, for whole cell voltage-clamp recordings from layer 2-3 pyramidal neurons. Because sleep-loss is associated with an increased extracellular concentration of adenosine throughout the forebrain (Porkka-Heiskanen, Strecker, & McCarley, 2000; Porkka-Heiskanen et al., 1997) which decreases glutamatergic mEPSC frequency and amplitude (Brambilla, Chapman, & Greene, 2005; Scanziani, Capogna, Gahwiler, & Thompson, 1992), all recordings were made in the presence of adenosine receptor 1 and 2 blockade (cyclopentyl-theophyline, 1μM).

As expected, we observed an increase of mEPSC frequency and amplitude after SD and the recovery to CS-levels in RS condition in the WT and *Mef2c^f/f^* mice (Fig. 4 A-F; Fig. S5; Table S6).

**Fig. 4.**
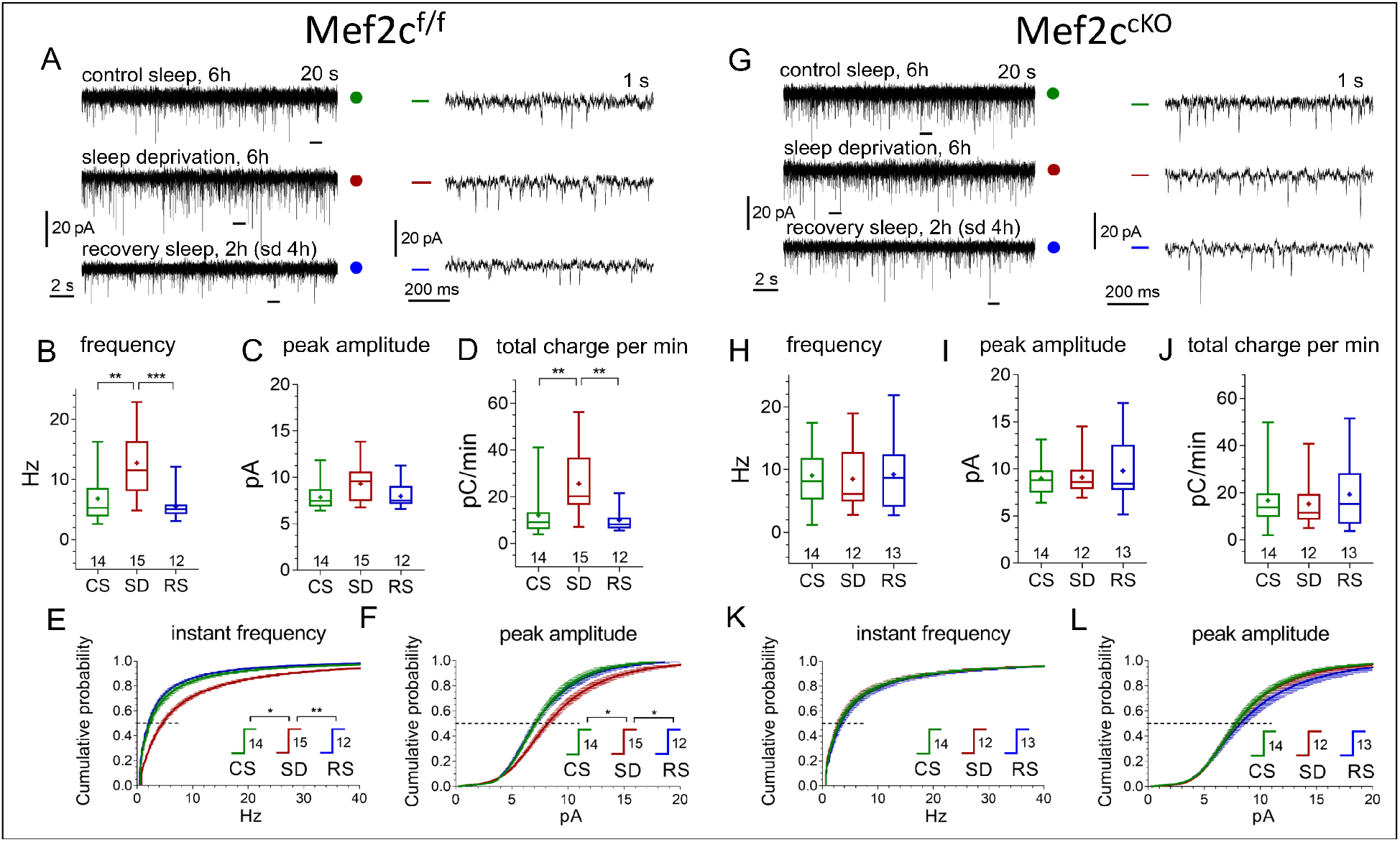
Conditional Mef2c knock-out in forebrain excitatory neurons eliminates sleep/wake remodeling of synaptic excitatory inputs to pyramidal neurons in anterior cingulate cortex slices. **(A, G)** Representative recordings of miniature excitatory postsynaptic current (mEPSC) traces (20 s, left panels, and expanded 1s traces on the right corresponding to time bars underneath original traces) obtained for three different experimental sleep/wake conditions, CS (green), SD (red), and RS (blue), for *Mef2c^f/f^* mice **(A)** and *Mef2c^cKO^* mice **(G)**. (**B,C,D**) and **(H,I,J**) illustrate mEPSC functional parameters obtained from *Mef2c^f/f^* and *Mef2c^cKO^* (number of cells for each condition shown above X axis condition label; also see Fig. S5 for comparison to WT). Averaged cumulative probability histograms for instant frequency and peak amplitude show increased frequency and amplitude with SD compared to CS or RS for *Mef2c^f/f^* (**E, F**) but not for *Mef2c^cKO^* mice (**K, L**). See methods and Table S5 for details and statistics.

To determine if the SD-induced increase in mEPSC frequency was caused by an increased presynaptic release probability or an increase in synapse number, we examined the paired-pulse (P_2_/P_1_) ratio from CS to SD and RS (Fig. 5 A, B).

**Fig. 5.**
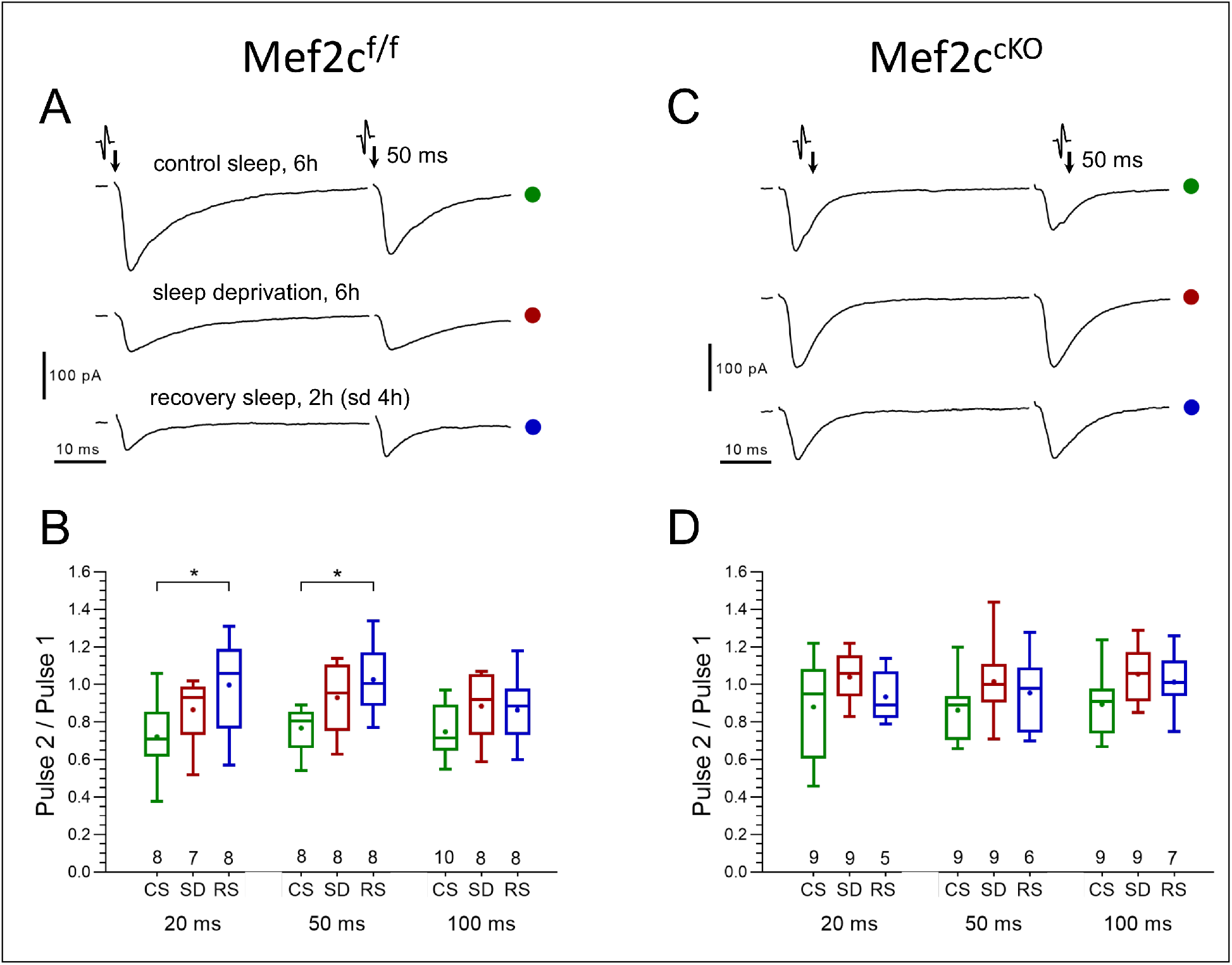
No significant differences in presynaptic glutamate release probability between CS and SD are found in either control or Mef2c^cKO^ mice. (**A** and **C**) Representative paired-pulse recordings of excitatory postsynaptic currents (evoked EPSC) evoked in layer 2/3 pyramidal neurons by two brief electrical stimulations of axon terminals in layer I, 50 ms interval, obtained for three different experimental sleep/wake conditions (CS, SD, RS) imposed on *Mef2c^f/f^* mice (**A**) and *Mef2c^cKO^* mice (**C**) respectively. The probability of presynaptic release (P_release_), is inversely proportional to the ratio of EPSC amplitudes (1/P_release_ ~ P_2_/P_1_); The P_2_/P_1_ ratio, obtained for three different experimental sleep/wake conditions (CS, SD, RS) and calculated separately for three different inter-pulse intervals (20, 50 and 100 ms), in *Mef2c^f/f^* (**B**) vs. *Mef2c^cKO^* mice (**D**). The plots (see Methods, numbers of experiments for each sleep/wake conditions shown underneath boxes), show no significant differences between CS and SD conditions in either control or in *Mef2c^cKO^* mice.

The P_2_/P_1_ ratio tended towards an increase from CS to SD condition (this is consistent with a decreased probability of release), suggesting that the SD-associated increase in mEPSC frequency resulted from an increased number of active synapses. However, for CS to RS, the observed significant increase in P_2_ to P_1_ ratio and the normalization of mEPSC frequency after RS is due to either a decrease in release probability, a decrease in active synapse number or both.

In *Mef2c^cKO^* mice, mEPSC frequency and amplitude was greater than in *Mef2c^f/f^* in CS condition. Consistent with the lack of change in the Mef2c^cKO^ transcriptome, a significant change in frequency and mEPSC amplitude in response to SD and RS sleep conditions was absent (Fig. 4 G-L, Table S7). Finally, as expected from the absence of SD or RS induced effects on mEPSCs, the *Mef2c^cKO^* mice had no change in P_2_/P_1_ ratio across sleep conditions (Fig. 5 C, D). These observations support a necessary role for *Mef2c* in SD and RS associated changes in synaptic strength for FC layer 2/3 excitatory neurotransmission.

**Fig. 6.**
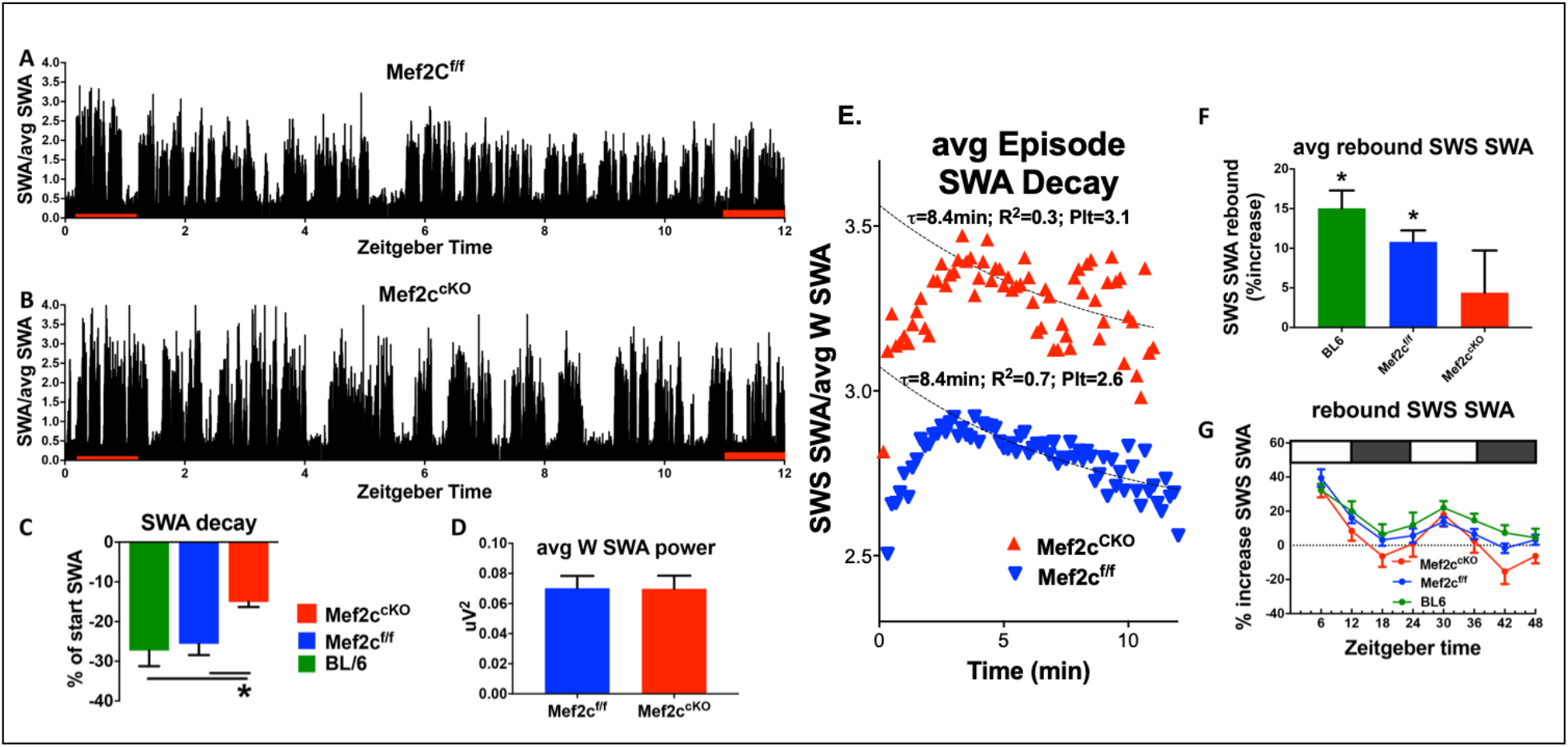
Loss of *Mef2c* increases sleep need and decreases its resolution. (**A**) SWA power (normalized to average SWA power over 24h) declines over the light phase as shown in a *Mef2c^f/f^* mouse. (**B**) The decline is less apparent in an example from a *Mef2c^cKO^* mouse. (**C**) Pooled data (*Mef2c^f/f^*, n=6; *Mef2c^cKO^*, n=11; C57BL/6, n=6) SWA power decay over the light phase for WT (BL/6), *Mef2c^f/f^* and *Mef2c^cKO^*. (**D**) The average waking SWA power in 24h is similar for *Mef2c^cKO^* and *Mef2c^f/f^* and is used as a normalizing factor to assess averaged SWS episode power. (**E**) SWS-SWA power over time from an average of SWS episodes aligned from the time of transition from waking to SWS, set to t=0. The time constant of decay, τ= 8.4 minutes, was calculated by best fit of the decay of power for the *Mef2c^f/f^* but the decay fit was indeterminate for *Mef2c^cKO^* (see Methods and Materials section for details). By setting τ= 8.4 minutes, a best fit to single exponential decay provided a plateau value (minimal value), an extrapolated peak value and an indication of goodness of fit in comparison to Mef2c^f/f^. The SWS-SWA power for *Mef2c^cKO^* mice is greater than *Mef2c^f/f^* as determined by the plateau value (Plt) and the peak value. The quality of the fit is reduced as indicated by the R^2^ value. (**F**) The average SWS-SWA power of each genotype following repeated, regular bouts of 4 h SD, was significantly increased compared to SWS-SWA power under baseline conditions. *Mef2c^cKO^* mice failed to show this rebound response to sleep loss. (**G**) The time course of rebound across the 8 × 4 h SD periods indicates that the lack of rebound in *Mef2c^cKO^* is particularly prominent during the dark phase.

#### The role of *Mef2c* in the generation and resolution of sleep need

Sleep need, inferred from the duration of time in waking without sleep (Borbely, 1982; Franken et al., 2001), is directly correlated with the ensuing SWS-SWA power calculated from surface EEG recorded from control *Mef2c^f/f^* and *Mef2c^cKO^* mice. In WT and *Mef2c^f/f^* mice, SWS-SWA is greatest at the start of the sleep phase and resolves over the sleep phase course as the mice spend time in SWS (ZT 0-12 h; Fig. 6 A, C, E). In contrast, *Mef2c^cKO^* mice express, for their averaged SWS-SWA episodes, higher SWA (Fig. 6 E) that fails to normally resolve over the course of the sleep phase (Fig. 6 B, C). Another indicator of increased sleep need is increased consolidation of SWS as suggested by a rightward shift in the cumulative histogram of SWS episode duration going from CS to SD conditions (Bjorness et al., 2016), which is occluded in *Mef2c^cKO^mice* (Fig. S6 A).

With respect to arousal-state duration (waking vs SWS vs REM), *Mef2c^cKO^* mice had no apparent significant differences and there were minimal differences in the power distribution across the frequency spectrum (Fig. S6 B, C). In response to SD, the *Mef2c^cKO^* mice show an attenuated or absent rebound of SWS-SWA (Fig. 6 F, G). These observations suggest that a loss of MEF2C function both increases the need for sleep and attenuates its resolution during SWS.

A syndromic form of autism, MEF2C Haploinsufficiency Syndrome (MCHS), is associated with missense mutations that disrupt DNA binding of MEF2C and with sleep-disruption (Harrington et al., 2020; Paciorkowski et al., 2013). We examined constitutive gene dosage on SWS-SWA power of a global *Mef2c* heterozygous gene-deletion mouse model (*Mef2c^+/-^*) of MCHS under CS and SD conditions (Fig. S7). Under CS conditions, SWS-SWA power resolved across the sleep phase, unlike the *Mef2c^cKO^* mice, indicative of a resolution of sleep need. Similarly, *Mef2c^+/-^* mice showed resolution of sleep need over an average SWS episode, again, unlike *Mef2c^cKO^* mice, however, the rate of resolution (as indicated by the rate of decay of SWA in an average SWS episode) was slowed. On the other hand, SD did not induce the expected rebound SWS-SWA in *Mef2c^+/-^* mice as was also the case with *Mef2c^cKO^* mice. Another similarity of *Mef2c^+/-^* mice with the *Mef2c^cKO^* mice is the apparent occlusion of the rightward shift in the cumulative histogram of SWS episode duration going from CS to SD conditions (Fig. S8), associated with increased sleep need in WT mice (Bjorness et al., 2016). The *Mef2c^+/-^* mice are already shifted to the right compared to *Mef2c^+/+^* mice in both CS and SD conditions. Furthermore, the *Mef2c^+/-^* mice had an abnormal relative increase in waking SWA power (in the high delta/low theta range of 4-5 Hz; Fig. S8).

By comparing differential gene expression under CS conditions in each of the *Mef2c* models, we found a significant overlap of genes changing in the *Mef2c^+/-^* and *Mef2c^cKO^* compared to their respective controls (142 overlapping genes; p-value =5.28e-08, Fisher’s exact test (Fig. S9, Table S8)). The identified overlapping gene set can provide clues into the molecular mechanisms underlying a requisite *Mef2c* sufficiency. Taken together, these findings suggest constitutive loss of one *Mef2c* allele is sufficient to alter the normal response to sleep loss but the loss appears to be at least partially compensated when spontaneous sleep remains undisturbed. More acute, complete loss of *Mef2c* function even further disrupts the homeostatic sleep loss response.

## Discussion

Sleep-loss induced genomic alterations are remarkably extensive, involving almost half of the expressed genome of the frontal cortex. About two thirds of the significant changes in gene expression observed in response to sleep deprivation were maintained when two hours of recovery sleep followed four hours of sleep loss indicating that direct recovery of sleep deprivation-induced transcriptional changes is delayed. This result suggests additional post-transcriptional mechanisms are also key to sleep-loss molecular mechanisms as for example, in synaptosomes, the translation of DEGs to proteins is at least, in part, dependent on sleep (Noya et al., 2019) or, more generally, the sleep loss associated increase in phosphorylation of the proteome (Wang et al., 2018).

Based on the shared transcriptional changes with sleep deprivation and recovery, we noted relevant increased expression of genes involved in protein anabolism (including genes encoding ribosomal components and enzymes controlling their activity), protein catabolism and mobilization of energy resources (including genes encoding mitochondrial structural components and enzymes). These altered expression patterns are associated especially with non-neuronal brain tissue. In contrast, there is enrichment for decreased expression of genes primarily linked to neuronal cells, for the control of synaptic strength. These sleep condition associations relate protein and energy metabolism to neuronal synaptic strength in much the same way as the energy-related metabolite, adenosine, mediates a synaptic homeostasis of glutamate synaptic activity (*Brambilla et al., 2005*), which is part of a neuronal-glial circuit limiting glutamatergic activity. This circuit is an essential part of the CNS response to sleep loss (*Bjorness et al., 2016; Bjorness, Kelly, Gao, Poffenberger, & Greene, 2009; Brambilla et al., 2005; Greene, Bjorness, & Suzuki, 2017*).

Coordination of the extensive change in transcriptome in response to loss of sleep requires the function of the transcription factor MEF2C. The MEF2C-dependent sleep loss DEGs encode proteins modulating cellular and systemic functions that are also expected to be affected. Indeed, the normal reduction of synaptic strength and activity associated with recovery sleep (RS) is absent in the *Mef2c^cKO^* mice. Notably, in WT, there is an acute increase in synaptic activity and strength induced by prolonged waking (SD) that we observe in association with a downregulation of genes controlling synaptic strength (Figs. 1, 4). These SD, down-regulated, synaptic DEGs are presumably not translated during the prolonged waking time, but with recovery sleep, their effects become manifest. Nonetheless, the normal increase in synaptic activity resulting from prolonged waking is also absent in *Mef2c^cKO^* mice, indicating an abnormal synaptic response both to the acute loss of sleep and to recovery sleep in the absence of MEF2C function. Even under *ad libitum* sleep conditions (i.e. in CS) loss of MEF2C function increased synaptic strength; however, this may be in response to the more chronic absence of MEF2C-mediated response to sleep loss.

Our findings show that loss or reduction of MEF2C function alters the response to sleep loss. Postnatal loss of MEF2C in forebrain excitatory neurons results in an enhanced SWS-SWA rebound and an attenuated resolution of the rebound. To the extent that SWS-SWA reflects sleep need, loss of MEF2C function increases sleep need in response to a loss of sleep compared to wild type MEF2C response, and it attenuates the sleep-mediated resolution of sleep need.

Taken together, these findings suggest a model of the sleep loss response that reflects a MEF2C-dependent, metabolically driven non-neuron to neuron homeostatic sleep circuit. This circuit can act to promote protein and energy metabolism and constrain synaptic activity in response to sleep loss.

## Acknowledgments

We wish to thank Mr. To Thai, Ms. Lilian Zhan and Dr. Busra Goksu for technical assistance and Dr. Stefano Berto for discussions about genomic analyses.

## Funding

This study was funded by NIH grant R01 MH080297 and WPI program for IIIS to R.W.G.; NIH grants RO1 MH102603, DC014702 and the James S. McDonnell Foundation 21^st^ Century Science Initiative in Understanding Human Cognition – Scholar Award (220020467) to G.K.; NIH grants R01 AG045795, R01 NS 106657 to J.S.T.; J.S.T is an Investigator in the Howard Hughes Medical Institute; NIH grant RO1 MH111464 to CWC and F30 HD098893 to CMB The contents do not represent the views of the U.S. Department of Veterans Affairs or the United States Government.

## Author Contribution

Conception and overall design by RWG; AK and AS directed by GK with consultation from JST for genomics design and analysis; CB and CWC for antibody design and analysis of MEF2C expression and phosphorylation; AJH and CWC for *Mef2c^+/-^* mouse, VR for synaptic functional design, assessment and analysis; TEB for EEG/EMG design, assessment and analysis. TEB, AK, VR, CWC, GK and RWG contributed to writing of the manuscript. All authors significantly contributed to manuscript discussion and revision.

## Competing interests

Authors declare no competing interests.

## Data and materials availability

The NCBI Gene Expression Omnibus (GEO) accession number for the RNA-seq data reported in this paper is GSE 144957.

## Supplementary Materials

### Materials and Methods

#### Animals

All mice (*Mus musculus*) had a C57BL/6 background, were male and of 8-19 weeks of age at time of data collection. Mice were on a 12 hour light-dark cycle with access to food and water ad libitum. For RNAseq experiments, we employed three genotypes: wild type (WT); a knock-in of loxP sites flanking exon 11 of the *Mef2c* gene, the second coding exon for the DNA binding site (Arnold et al., 2007)(*Mef2c^f/f^*), a control for the *Mef2c^cKO^;* and a CamKII-promoter (Tsien et al., 1996), CRE-mediated recombination of the *Mef2c^f/f^* loxP sites (founders obtained from Jackson Labs Jackson labs; B6.Cg-Tg(Camk2a-cre)T29-1Stl/J), resulting in a region and cell type specific conditional knockout (*Mef2c^cKO^* (Arnold et al., 2007)). Primers and primer products are shown along with a map of the *Mef2c* gene(Figure M1). Initially, we used a primer pair, Mef2C.2 and Mef2C.3 that gave a product of 650 BP for WT, 750BP for *Mef2c^f/f^* and no product for a CRE-mediated recombined allele to distinguish homozygous and heterozygous floxed knockins from WT mice.

**Fig. M1.**
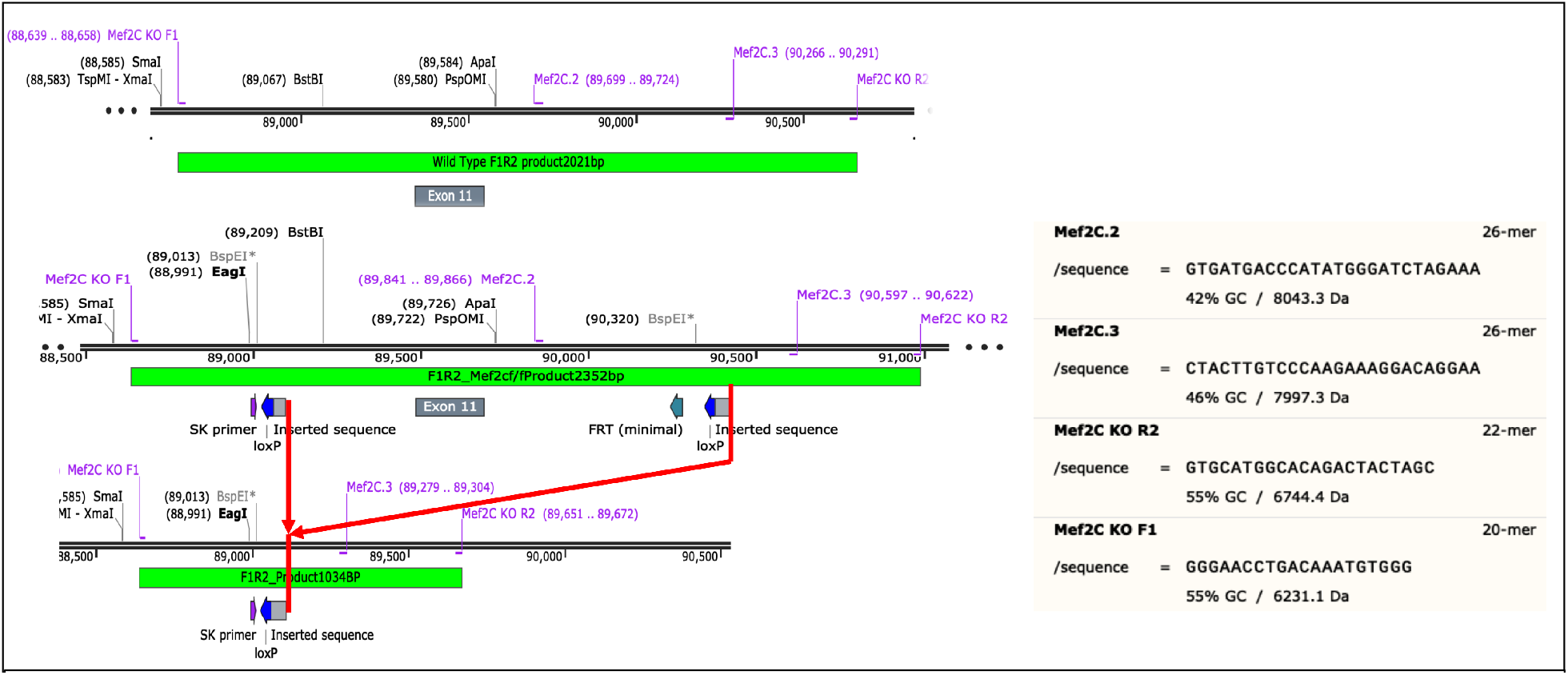
Map of *Mef2c* showing primer pairs and loxP sites for *WT, Mef2c^f/f^* and the recombined *Mef2c^cKO^* genes. Note that exon 11 is the second coding gene for *Mef2c*.

**Fig. M2.**
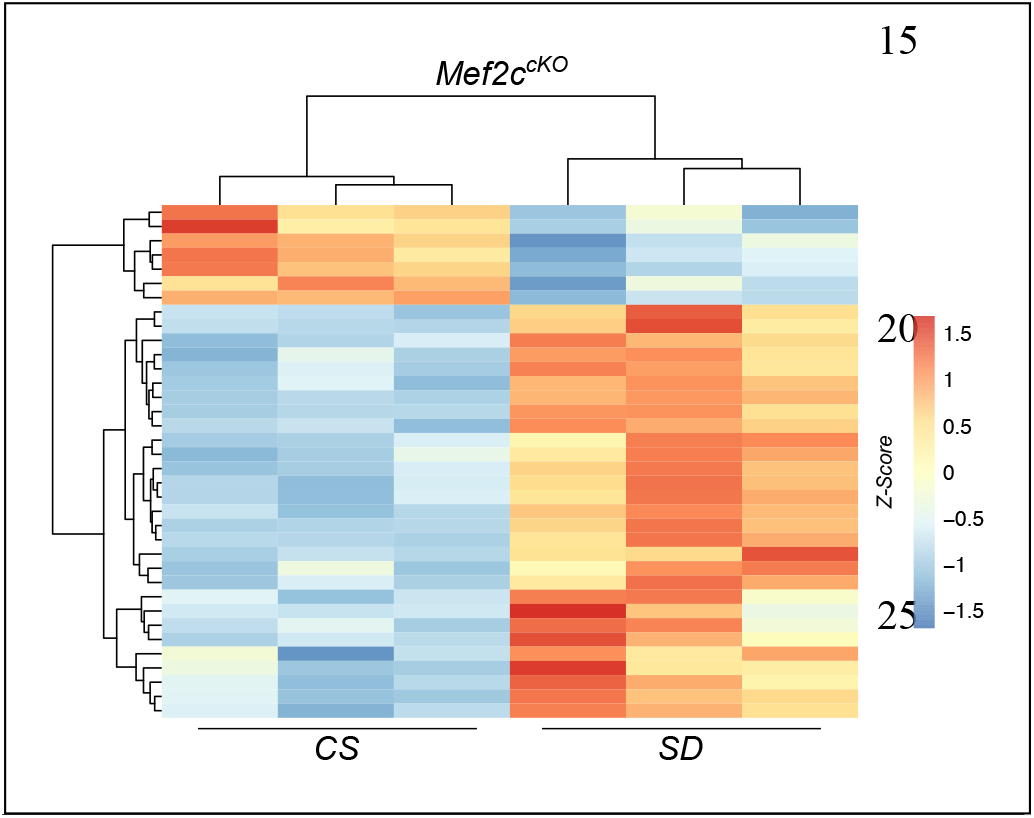
Heat map of DEGs from *Mef2c^cKO^* mouse samples under CS or SD conditions. The sample from the second column (CS) showed a similar DEG clustering pattern to the other two samples that had germline recombination.

In order to confirm CRE-mediated recombination in the brain, we used another primer pair, Mef2C KO F1 and Mef2C KO R2 (Fig. M1) that gave products of 2250 BP for non-recombined *Mef2c^f/f^* and 1000 BP for recombined *Mef2c^cKO^* alleles. All brain samples of *Mef2c^cKO^* mice expressed both products as expected.

We also determined that some of the offspring of male *Mef2c^f/wt^;CamKII:Cre* mice were heterozygote for a single recombined allele, showing *Mef2c^f/-^* in tail samples, consistent with CRE-mediated recombination in spermatocytes (Luo et al., 2020). Thus, genedosage dependent developmental effects of *Mef2c* loss of function in all cell types may contribute to the transcriptome of *Mef2c^cKO^* mice in addition to the homozygous conditional loss of *Mef2c* function in the brain. Nevertheless, a heat map of differentially expressed genes (DEGs) from *Mef2c^cKO^* mice showed similar clustering of DEGs from samples in CS conditions with, and without (second column Fig. M2) recombination in the tail.

For in vivo electrophysiology (EEG/EMG analysis for sleep/waking state determination), in addition to the three strains described above, we also included constitutive heterozygous Mef2c knockouts (*Mef2c^+/-^*) and their littermate controls (*Mef2c^+/+^);* generation of these strains has been described previously (Harrington et al., 2020).

All experimental procedures were approved by the University of Texas Southwestern IACUC or VA North Texas Health Care System IACUC.

#### Sleep Deprivation Protocol

All mice underwent an accommodation protocol of 3 weeks duration, in 12/12 light/dark schedule, in plexiglass, false-bottomed cages, suspended over a treadmill with ad libitum water and mouse chow as previously described (Bjorness et al., 2016). The treadmill (speed of treadmill = 3cm/sec) was turned on daily for 10 minutes to accommodate the mice to the slow movement (mice are able to eat and drink with it running) and to clean it. For RNAseq, 36 hours prior to the experimental day (CS, SD or RS) a dark/dark schedule was instituted. The use of the treadmill prevented transition to sleep, did not increase motor activity (due to the slow speed) and provided a mild, familiar, non-varying, arousing stimuli that reduced the experimental variability associated with more traditional “gentle handling” techniques.

#### RNA Collection, RNA-seq library preparation and Sequencing

Total RNA was extracted from all the samples using TRIzol reagent (Thermo Fisher Scientific, Cat. No. 15596026) and was purified with RNeasy mini kit (Qiagen, Cat No. 74104) according to manufacturer’s instructions. All the samples were then randomized in order to reduce the batch effect and submitted to McDermott Sequencing Core at University of Texas Southwestern Medical Center for library preparation and sequencing. Libraries were prepared using TrueSeq Stranded mRNA Library Prep (Illumina, Cat. No. 20020596) as per manufacturer’s instructions by McDermott Sequencing Core. The quality and concentration of the libraries was checked on Agilent Bioanalyzer High Sensitivity DNA chip (Agilent, Cat. No. 5067-1504). Samples were pooled and sequenced by McDermott Sequencing Core using Illumina’s NextSeq500 to yield paired-end (150bp long) strand-specific reads.

#### RNA-seq Data Analysis

De-multiplexed raw reads were received from McDermott Sequencing Core. Raw reads were first filtered for phred quality and adapters using FASTQC (*FastQC, Babraham Bioinformatics, URL:* https://www.bioinformatics.babraham.ac.uk/projects/fastqc) and Trimmomatic. Filtered reads were then aligned to the reference mouse genome mm10 (https://genome.ucsc.edu) using STAR (Dobin et al., 2013) aligner. Uniquely mapped reads were used to obtain the gene counts using HTSeq package (Anders, Pyl, & Huber, 2015) and the read counts were normalized using the CPM (counts per million) method implemented in the edgeR package (McCarthy, Chen, & Smyth, 2012; Robinson, McCarthy, & Smyth, 2010). For further analysis, we performed a sample-specific CPM filtering, considering genes with CPM values of 1.0 or greater in all replicates in one or more of the sleep conditions. DESeq2 (Anders & Huber, 2010; Love, Huber, & Anders, 2014) was used to detect the differentially expressed genes (DEGs) across sleep states. We applied a filter of an adjusted *p*-value of ≤ 0.05 and absolute log fold change of ≥ 0.3 to identify DEGs. Significant DEGs are visualized using volcano plots. *UpSetR* (Conway, Lex, & Gehlenborg, 2017) R package was used to identify shared and unique sets of DEGs across sleep states.

#### GO Analysis

GO analysis of the significant, sleep-states shared and unique DEGs was performed using *ToppGene (*https://toppgene.cchmc.org) and GO terms were reduced using *REVIGO* (Supek, Bosnjak, Skunca, & Smuc, 2011) GO categories were considered significant if they contained at least three genes and if they had a Benjamini–Hochberg (B–H)-corrected *p*-value of ≤ 0.05.

#### DEG enrichment against cell types, MEF2C targets and rPRGs

The mouse visual cortex single cell RNA-seq dataset was downloaded from NCBI Gene Expression Omnibus (accession no. GSE102827) and processed using *Seurat* (Butler, Hoffman, Smibert, Papalexi, & Satija, 2018). Retaining cluster annotations from the original source (Hrvatin et al., 2018), genes enriched in each cell-type/cluster were identified using the *FindAllMarkers* function of *Seurat* (Butler et al., 2018) with default parameters. The list of MEF2C target genes was obtained from (Harrington et al., 2016). The list of rapid primary response genes was obtained from (Tyssowski et al., 2018). The list of differentially expressed genes from *Mef2c^+/-^* compared to *Mef2c^+/+^* at CS was obtained from (Harrington et al., 2020). Enrichment analyses for shared and unique list of DEGs across sleep states against (i) cell types from mouse visual cortex single cell datasets, (ii) MEF2C targets and (iii) rPRGs were performed using a Fisher’s exact test. Also, enrichment analysis for differentially expressed genes across *Mef2c^f/f^* and *Mef2c^cKO^* under CS and differentially expressed genes from *Mef2c^+/-^* compared to *Mef2c^+/+^* at CS was performed using a Fisher’s exact test.

#### Weighted Gene Co-expression Network Analysis

Weighted gene co-expression network analysis (WGCNA) was performed on 19 total RNA-seq samples (6 control sleep (CS) samples, 6 sleep deprived (SD) and 7 recovery sleep (RS) samples. The R package for WGCNA (Langfelder & Horvath, 2008) was used to build a gene co-expression network using filtered CPM data (CPM > = 1 across all replicates of one or more of each of the conditions). A signed network was constructed using the *blockwiseModules* function of the WGCNA R package. A value of 26 was chosen as Beta with highest scale-free R square (R^2^ = 0.796). For other parameters, we used *corType = pearson, maxBlockSize = 15000, reassignThreshold = 1×10^−5^, mergeCutHeight = 0.05, deepSplit = 4, detectCutHeight = 0.999* and *minModuleSize = 50*. Visualizations of network plots were created using Cytoscape v3.4.0(Shannon et al., 2003), representing the top 250 edges based on ranked weights.

#### Heat map plot for enrichment of gene lists against WGCNA module genes

List of genes per WGCNA module were enriched against (i) DEGs per sleep state, (ii) cell type genes identified in mouse cortex (Hrvatin et al., 2018), (iii) MEF2C targets (Harrington et al., 2016), and (iv) module eigengene correlation to each sleep state using *supertest* function of *SuperExactTest* R package. The heat map is generated using *ggplot2* functions *geom_tile*. The modules are ordered based on module eigengene correlation to CS sleep state starting from highest to lowest correlation.

#### Immunoprecipitation of MEF2C Protein

Frozen brain tissue from the frontal cortex of mice were immediately homogenized by sonication after addition of 600 μL mRIPA buffer with inhibitors (50 mM Tris-HCl pH 7.5, 150 mM NaCl, 1mM EDTA, 1% iGEPAL CA-630, 0.5% sodium deoxycholate, 0.1% SDS, 200 mM sodium fluoride, 1 mM cyclosporine A, 200 mM sodium orthovanadate, 1 pellet of complete, EDTA-free tablets (Sigma/Roche 11836170001) in deionized water for 10 mL total volume of solution). Samples were then centrifuged at max speed and supernatant was aliquoted. Protein concentration was quantified via Bio-Rad DC Protein Assay (Bio-Rad 5000114). Protein lysates (1.5 mg) were diluted to a total volume of 500 μL. A volume of 2 μL of mouse monoclonal anti-MEF2C antibody (Novus NBP2-00493) was added to each diluted protein lysate and incubated for 2 hours at 4°C. Samples were then added to washed protein G Plus/A agarose beads (Millpore IP05) and incubated for 1 hour at 4°C. The samples were then washed three times with mRIPA buffer with inhibitors and eluted with 2X sample buffer + 10% 2-mercaptoethanol.

#### Western Blot of MEF2C Immunoprecipitation

Immunoprecitation (IP), pre-IP, and post-IP samples were loaded into 7.5% Mini-PROTEAN TGX Stain-Free gels (Bio-Rad 4568026). Gel electrophoresis was run at 250 V for 25 minutes in 10X Tris-gycine SDS buffer. Proteins were then transferred onto PVDF membranes using the Bio-Rad Trans-Blot Turbo System on the high MW setting. Membranes were blocked in 1:1 Odyssey blocking buffer/1X PBS for 1 hour. Membranes were incubated overnight with primary antibodies: rabbit anti-phospho-MEF2 (1:200, lab purified) or rabbit anti-MEF2C (1:2500, Abcam ab197070). After washing, Licor secondary antibodies were incubated on the membranes (1:20,000, Licor IRDye 800CW goat anti-rabbit 926-32211). After washing, blots were imaged on the Licor Odsussey CLx system. The pre-IP sample membranes were then incubated with rabbit anti-beta-actin antibody overnight. After washing, the membrane was incubated with the Licor goat anti-rabbit secondary and imaged.

#### Analysis of MEF2C Immunoprecipitation

Licor image files were imported into Licor ImageStudio software and protein bands were quantified. We calculated a ratio for phospho-MEF2C signal to the total MEF2C signal for each sample. The average ratio for the control samples was used as a normalization factor for each sample. The ratio for each sample were then divided by the normalization factor to generate normalized signal values for each sample. Similar analysis was used to compare total MEF2C signal to beta-actin signal. One-way ANOVA statistical analyses were performed in Graphpad Prism software.

#### Brain slice electrophysiological recordings

Brain slices 300 μm thick, cut 15°-angled in rostral direction to keep perpendicular blade penetration in the anterior cingulate area, were prepared from 8-12 week old control (Mef2c fl/fl: Cre-) and Mef2c KO (Mef2c fl/fl: Cre+) mice at the same Circadian time ZT6 immediately after subjecting to one of the three 6-hour sleep/wake protocols (CS, SD, RS). The dissection was done in ice-cold solution containing (mM): NaCl 84, KCl 3, NaH_2_PO_4_ 1.25, CaCl_2_ 0.5, MgSO_4_ 7, NaHCO_3_ 26, glucose 20, sucrose 70, ascorbic acid 1.3, kynurenic acid 2 (brought to pH 7.3 by continuous saturation with carbogen gas containing 95% of O_2_ and 5% of CO_2_). After dissection, slices were transferred to the carbogen gas-saturated artificial cerebrospinal fluid (ACSF) solution containing (mM): NaCl 125, KCl 5, NaH_2_PO_4_ 1.25, CaCl_2_ 2, MgCl_2_ 1.3, NaHCO_3_ 25, glucose 12 (pH 7.3) at 32° C, and were used for experiments after 1 hour equilibration.

Slices were placed in 400 μl recording chamber and continuously perfused at 2 ml/min with heated (33° C) carbogen gas-saturated ACSF containing 1 μm of adenosine receptor 1 blocker 8-cyclopentyl theophylline (CPT) during entire experiment. Solutions containing additionally 100 μm of picrotoxin (PTX) or 1 μl tetrodotoxin (TTX) were washed-in to block GABA receptor synaptic currents during paired-pulse stimulation experiments, and Na^+^ voltage-dependent channel-generated spontaneous presynaptic spikes during recordings of miniature excitatory post-synaptic currents (mEPSC) respectively.

Pyramidal neurons in cortical layers 2/3 at the anterior cingulate cortex (ACC) area were visually identified with upright microscope equipped with infra-red light sensitive camera, and voltage-clamped at −70 mV holding potential (Cl^−^ equilibrium potential in our solutions) with patch pipette containing (mM): K-methanesulfonate 130, NaCl 5, MgCl_2_ 2, Hepes 10, EGTA 1, CaCl_2_ 0.4 (free Ca^2+^ ~80 nM), MgATP 2, Na_2_GTP 0.2 (pH~7.3 adjusted with NaOH). Multiclamp 700A amplifier, Digidata 1440A and pClamp 10.2 software (all Molecular Devices, CA) were used for patch-clamp recordings both in voltage-clamp and current clamp modes. The identity of each pyramidal neuron was confirmed by recording a specific pattern of trains of action potentials in current-clamp mode, clearly different from fast-spiking activity of interneurons in response to a series of depolarizing current steps.

CaCl_2_ was removed, and 0.25 mM of QX 314, a Na^+^ voltage-dependent channel blocker was added to the patch pipette solution to suppress generation of postsynaptic Na^+^ currents during recordings of evoked excitatory post-synaptic current (eEPSC) in paired-pulse stimulation experiments. In these experiments, the identity of pyramidal neurons was verified immediately after breaking the membrane patch before suppression of action potentials. Stimulation of axons forming synapses at apical dendrites of L2/3 pyramidal cells during paired pulse experiments was done by placing a bipolar stimulation electrode in the L1 area at a distance ~ 100-200 μm from the recorded neuron axial projection to this area. Brief 0.4 ms biphasic stimulation current pulses of 30 to 60 μA delivered with Biphasic Stimulus Isolator BSI-950 (Dagan, MN) were enough to produce minimal-amplitude reproduceable eEPSCs in 95% of experiments.

#### Analysis of *mEPSC*

Functional parameters of miniature excitatory post-synaptic currents (mEPSCs) were calculated from 2 to 4 min continuous recordings following steady-state after application of 1 μl TTX. Recordings were high-pass filtered at 10 Hz and low-pass filtered at 500 Hz. Individual mEPSC were detected by the event template detection algorithm embedded into the pClamp 10 software. A combination of five mEPSCs templates varying from 6 to 25 ms durations was used, each template recognition window being “trained” from ~ 100 real events. No minimal threshold for mEPSC amplitude was applied for medium and longer events to avoid potential exclusion of low-amplitude attenuated mEPSCs originating from distal dendrites. The same template recognition set of parameters was applied for the analysis of data obtained from all cells included in this study. The microscopic parameter values of individual events (peak amplitude, area, instant frequency) were calculated by the PClamp event analysis algorithm and transferred to the Microsoft Excel for secondary off-line general statistics analysis. Standard box and whiskers graphs for each analyzed mEPSC parameter averaged for all cells within a particular experimental group were plotted using GraphPad Prism software, which represents 25% to 75% interval as vertical boxes, the whole range of data points as whiskers, the line inside boxes as a median, and a “+” sign inside boxes indicates the mean value. One-way ANOVA analysis with Tukey’s multiple comparisons test was used to define significant differences within three experimental groups (CS, SD, RS for each genotype) for each reported parameter.

Cumulative histograms for each measured mEPSC parameter of a particular experiment were genrated from a continuous 2 to 4 min recording and comprised values falling within a mean ± 3SD interval (99.7% of events). Within a particular experimental group (CS, SD, RS for each genotype), histograms from individual cells were binned at equal 0.01 steps and then averaged for each binning point over the entire Y-scale range (from 0 to 1). Averaged cumulative histograms were plotted using GraphPad Prism and included mean ± SEM (X-axis error bars) for each binning point. Ordinary one-way ANOVA analysis with Tukey’s multiple comparisons test was used to define significant difference between resultant averaged histograms within each genotype group.

##### Paired-pulse eEPSC analysis

For each paired-pulse experiment, peak amplitudes of coupled evoked EPSCs (eEPSCs) were obtained after averaging traces of 5 to 10 repetitive paired stimulation sequences separated by 20 s relaxation time. P2/P1 amplitude ratio was calculated for three interpulse intervals: 20, 50 and 100 ms. Standard box and whiskers graphs for each P2/P1 ratio averaged for all cells within a particular experimental group were plotted for each of three interpulse intervals using GraphPad

Prism software. Ordinary one-way ANOVA analysis with Tukey’s multiple comparisons test was used to define significant differences within three experimental groups (CS, SD, RS for each genotype) for each measured P2/P1 ratio.

#### EEG and EMG acquisition for sleep/waking state analysis

Animals (*Mef2c^cKO^* n=7, *Mef2c^f/f^* n=11, C57BL/6 n=6, *Mef2c^+/-^* n=6, *Mef2c^+/+^* n=6) were implanted with EEG and EMG electrodes using standard procedures (Bjorness et al., 2016). Briefly, mice were anesthetized with isoflurane and placed into a stereotaxic apparatus after which the hair is sheared and the scalp cleaned and incised. Four small holes were drilled for the placement of bilateral skull screw style electrodes over the frontal cortex (AP +1.7, ML +/- 1.77), parietal cortex (AP −1.7, ML +2.0), and occipital cortex (AP −5.5, ML −1.5). Paddle style EMG electrodes (Plastics One) were placed under the dorsal nuchal muscle. Electrode pins were gathered into a six pin pedestal (Plastics One) which was cemented to the skull using dental cement. The wound was closed using absorbable sutures and coated with triple antibiotic ointment. Mice received buprenorphine for analgesia and were given 14 days to recover from surgical procedures prior to experimental procedures.

Mice were placed into individual cages suspended above the belt of a custom treadmill apparatus with food and water available ad libitum. Mice were given two weeks to acclimate to the new environment with daily short duration exposure to the treadmill belt moving after which mice were connected to an amplifier system via a tether attached to the pedestal with an additional week of acclimation to the tether. A balance bar was set between the tether and commutator to allow for unrestricted movement. Following acclimation, EEG and EMG signals were acquired using a 15LT amplifier system (Natus Neurology) for two days under baseline (undisturbed) conditions followed by two days of chronic, partial sleep deprivation (Bjorness et al., 2016). Sleep deprivation consisted of eight cycles of four hours TM ON (belt moving) followed by two hours TM OFF (belt fixed). State assignment was determined via a custom Matlab (Mathworks) autosorter program using standard criteria (Bjorness et al., 2016) for waking, SWS, and REM after which files were manually checked to ensure correct assignment of state and flag epochs featuring artifact. Epochs were scored in 10 s bins with artifact-flagged epochs excluded from power spectral analysis. Episodes began with 30 consecutive seconds of one state and ended with 30 consecutive seconds of a different state. Power spectrum values were calculated in a 2 s window with a 1 s overlap and a Hamming window using the mean squared spectrum function in Matlab. For statistical analyses, *Mef2c^cKO^* was compared to *Mef2c^f/f^* and C57BL/6, while *Mef2c^+/-^* was compared to *Mef2c^+/+^*.

#### Decay of SWA across the light (inactive) phase

SWA (0.5-4.5 Hz) was normalized to the 24 h average SWA and the percentage change between the first and last consolidated 1 h sleep period was calculated as previously described (Nelson et al., 2013). The baseline day with the clearest consolidated sleep periods early and late in the light phase were used for statistical comparison via a Oneway ANOVA with Sidak’s multiple comparison test.

#### Decay of SWS-SWA within an averaged SWS episode under baseline conditions

For a full description of the decay analysis see (Bjorness et al., 2016). Briefly, SWS-SWA power was normalized to W SWA power after which SWS episodes of at least 5 min in duration were collected and gathered into an excel file with one episode per column such that the normalized SWS-SWA power of the first epoch was in the first row of each column, the normalized SWS-SWA power of the second epoch was in the second row of each column, and so forth. Based on the 5 min duration criteria, all columns had at least 30 rows; the longest 4 episodes were truncated to leave 5 episodes at the longest time point. One *Mef2c^cKO^*, one *Mef2c^f/f^*, and one *Mef2c^+/+^* mouse per group was excluded on the basis of insufficient episodes of at least 12 min in duration (Bjorness et al., 2016). The natural log was taken after which episodes were averaged within animal to create a single averaged SWS episode per mouse followed by averaging across animals within group to result in a single episode per group. Next, the group average episode was transformed by taking the exponential. A fragment of the average was created (excluding the initial ascending phase and periods of high variability towards the end) and fit using a single phase decay (GraphPad Prism) in order to determine the time constant of decay (⊤). Based on previously described exclusion criteria (absolute sum of squares, Bjorness et al., 2016), the averaged group episode for *Mef2c^cKO^* mice could not be fit. Thus, the time constant of decay determined for the *Mef2c^f/f^* group was used to determine the plateau and R^2^ value for the *Mef2c^cKO^* group.

#### Rebound SWS-SWA following chronic, partial SD

Mice underwent 8 cycles of 4h SD (TM ‘ON’) followed by 2h RS (TM ‘OFF’) starting at ZT0. SWS-SWA power was averaged in 2 h bins. Rebound SWS-SWA was determined by the percent change in SWS-SWA from baseline conditions to TM ‘OFF’ conditions using time-matched circadian bins (i.e. ZT 4-6, ZT 10-12, ZT 16-18, ZT 22-24). A Oneway ANOVA was used to compare average rebound across the 8 RS periods and a One sample T test was used to compare average values to a theoretical mean of 0 (GraphPad Prism).

#### Time in sleep/waking state

Time in each sleep/waking state (waking, SWS, REM) under baseline conditions was averaged in 2 h bins and calculated as the percent time of the total period. Averaged percent time in state for each state was compared between groups using a One-way ANOVA (GraphPad, Prism).

#### Cumulative distribution of SWS episodes

SWS episodes under baseline and SD (recovery periods) conditions were binned by duration using the histogram function in excel (Microsoft); the first bin was 90 sec with successive bins increasing by 40 sec. The number of episodes in each bin was divided by the total number of episodes for the condition and multiplied by 100 to get a percentage of the total episodes represented by each bin. Next, a cumulative percentage was calculated by adding the percentage of successive bins. A Two-Way ANOVA (condition, bin) was used to compare cumulative histogram episode durations across groups and conditions (baseline and SD).

#### Spectral power distribution across state

Under baseline conditions, individual 10s epochs were separated by state (waking, SWS, REM) after which spectral power in each 1 Hz bin up to 50 Hz was normalized by the total power for that state (waking epoch normalized to waking total power, etc). Fraction power was then averaged across epochs within each 1 Hz bin to get an average spectral distribution from 1 to 50 Hz by state. Two-way ANOVA (group, frequency) was used to compare spectral distributions by state across groups.

### Figures S1to S9

**Fig. S1.**
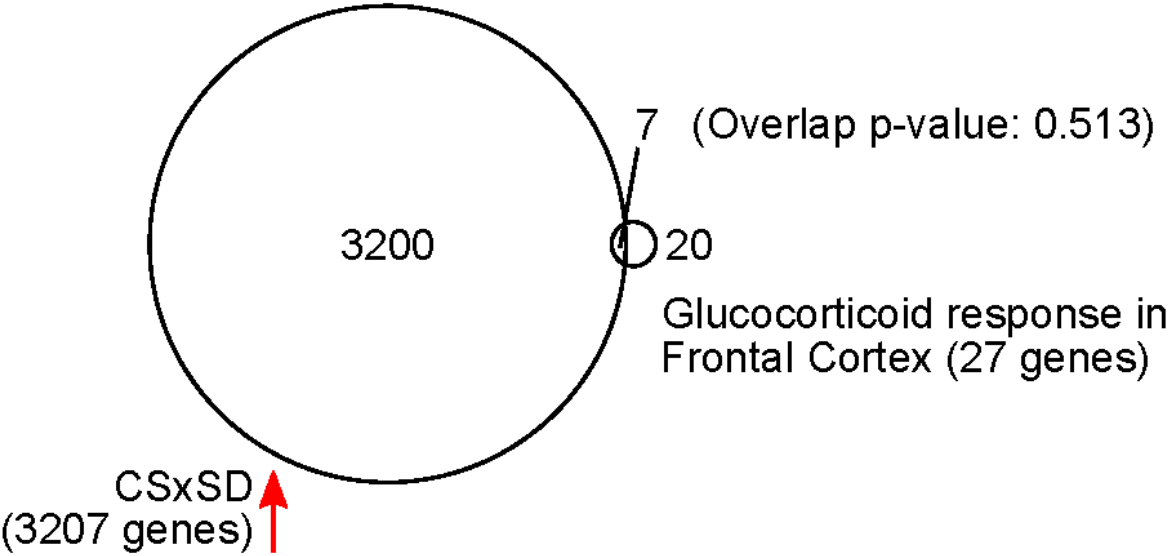
3207 genes increased their expression after six hours of sleep deprivation (SD). 27 genes are expressed in the frontal cortex and involved in glucocorticoid intracellular responses (MGI GO:0071385). Seven genes have both increased expression in response to SD and are involved in glucocorticoid responses (p-value=0.513 using Fisher’s exact test).

**Fig. S2.**
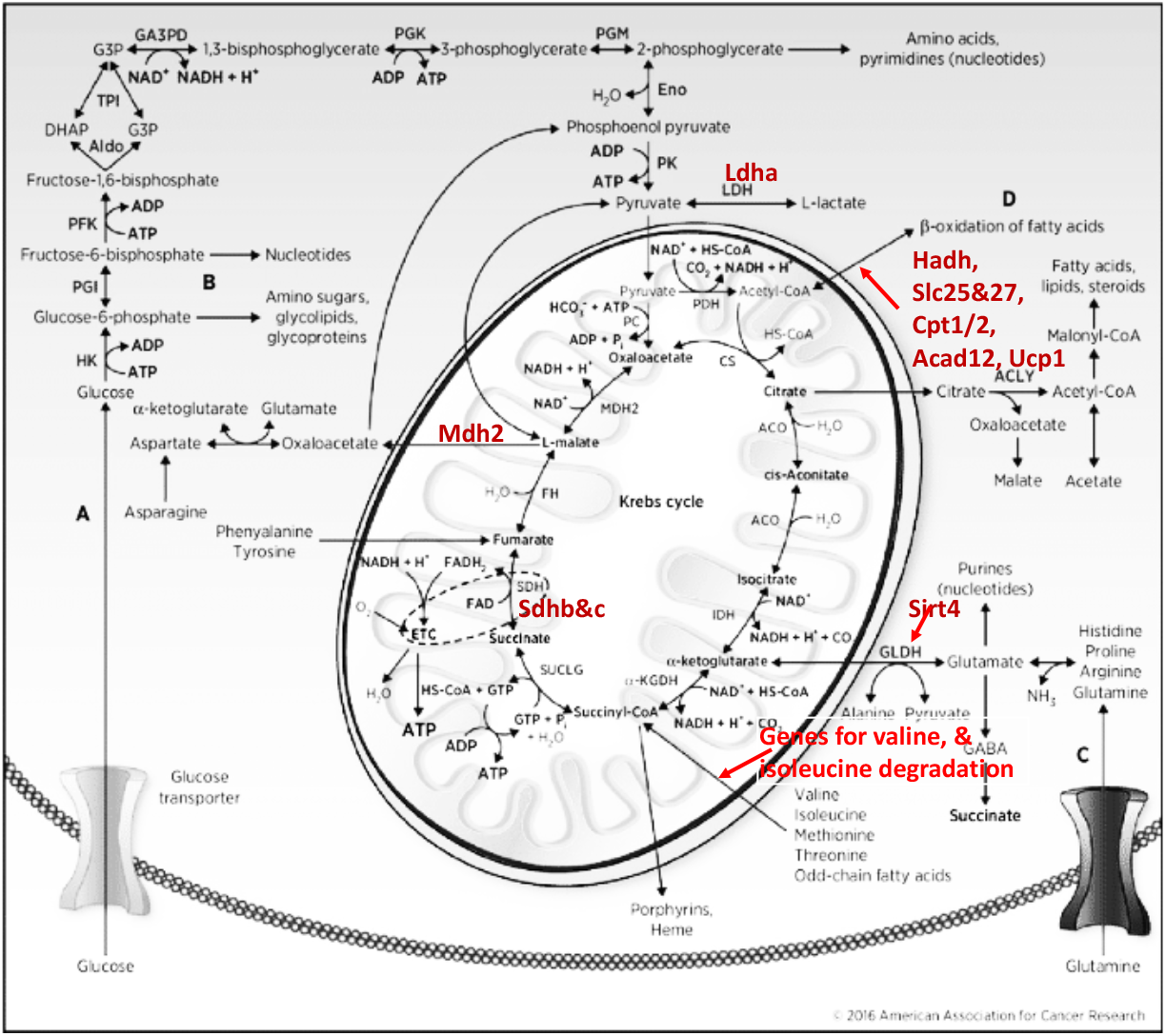
Kreb’s cycle with highlighted differentially expressed genes with increased expression in both SD and RS, and involved in anaplerotic reactions.

**Fig. S3.**
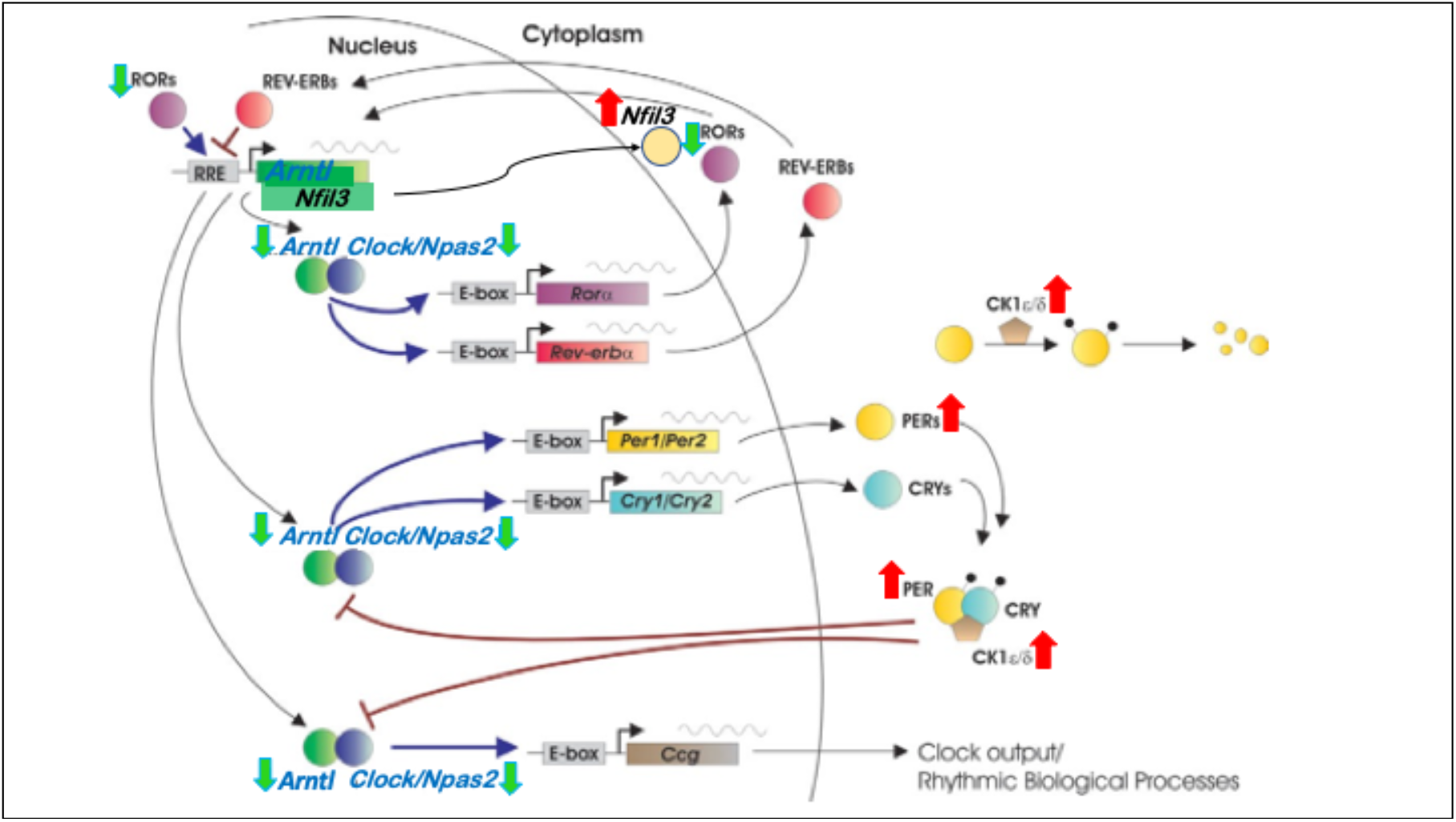
Circadian rhythm gene regulatory network with highlighted core clock genes that are affected during SD. Genes with increased expression in SD are marked with *red* up arrows and genes with decreased expression in SD are marked with *green* down arrows.

**Fig. S4.**
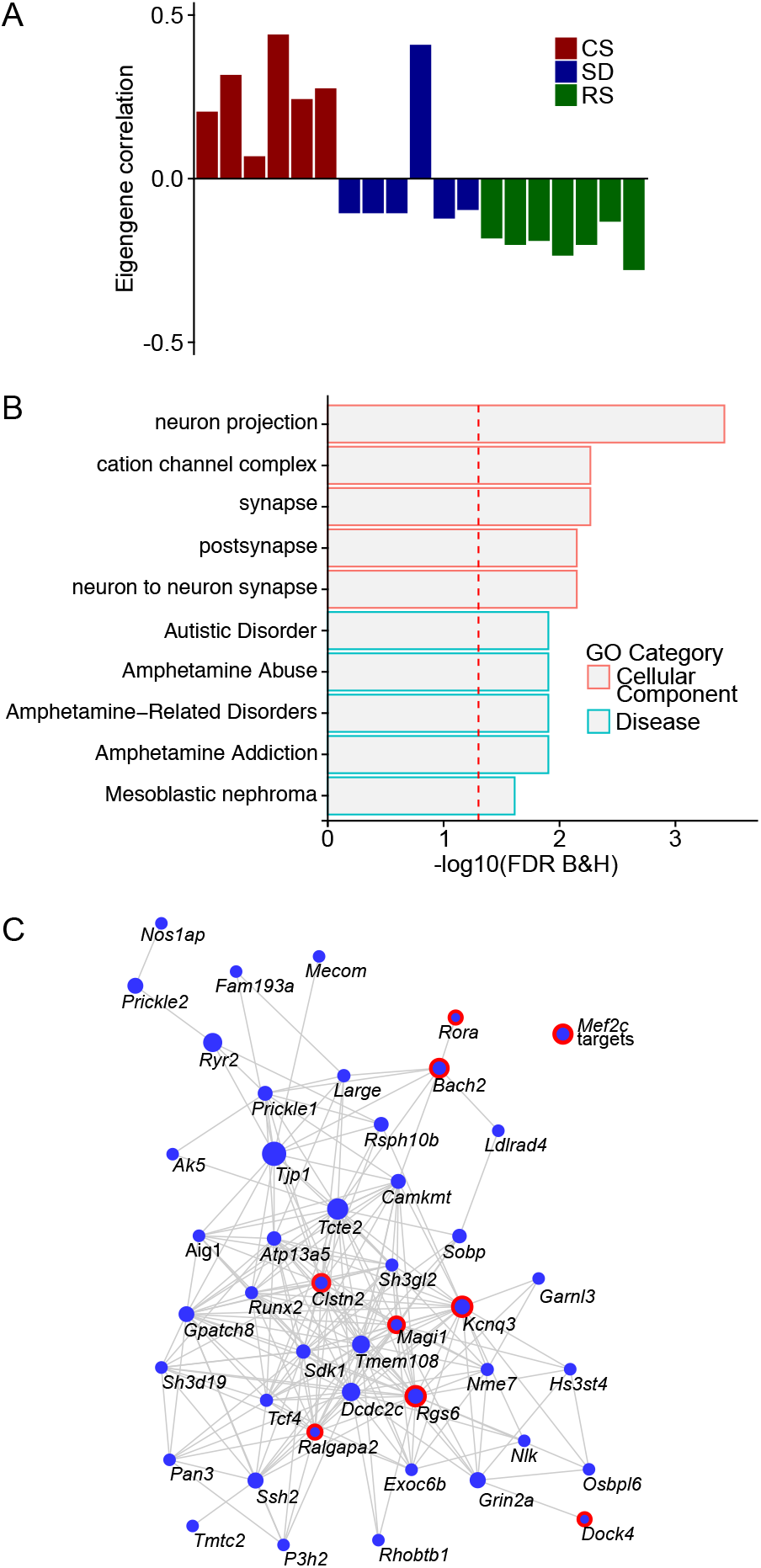
(**A**) Barplot for Module ID 10 with eigengene correlation across sleep states showing a strong positive correlation to CS and negative correlation to SD and RS. (**B**) GO plot for genes within the Module ID 10 (**C**) Network plot for Module ID 10 showing the top 250 connections between genes. Genes that are MEF2C targets (14) are highlighted by red circles.

**Fig. S5.**
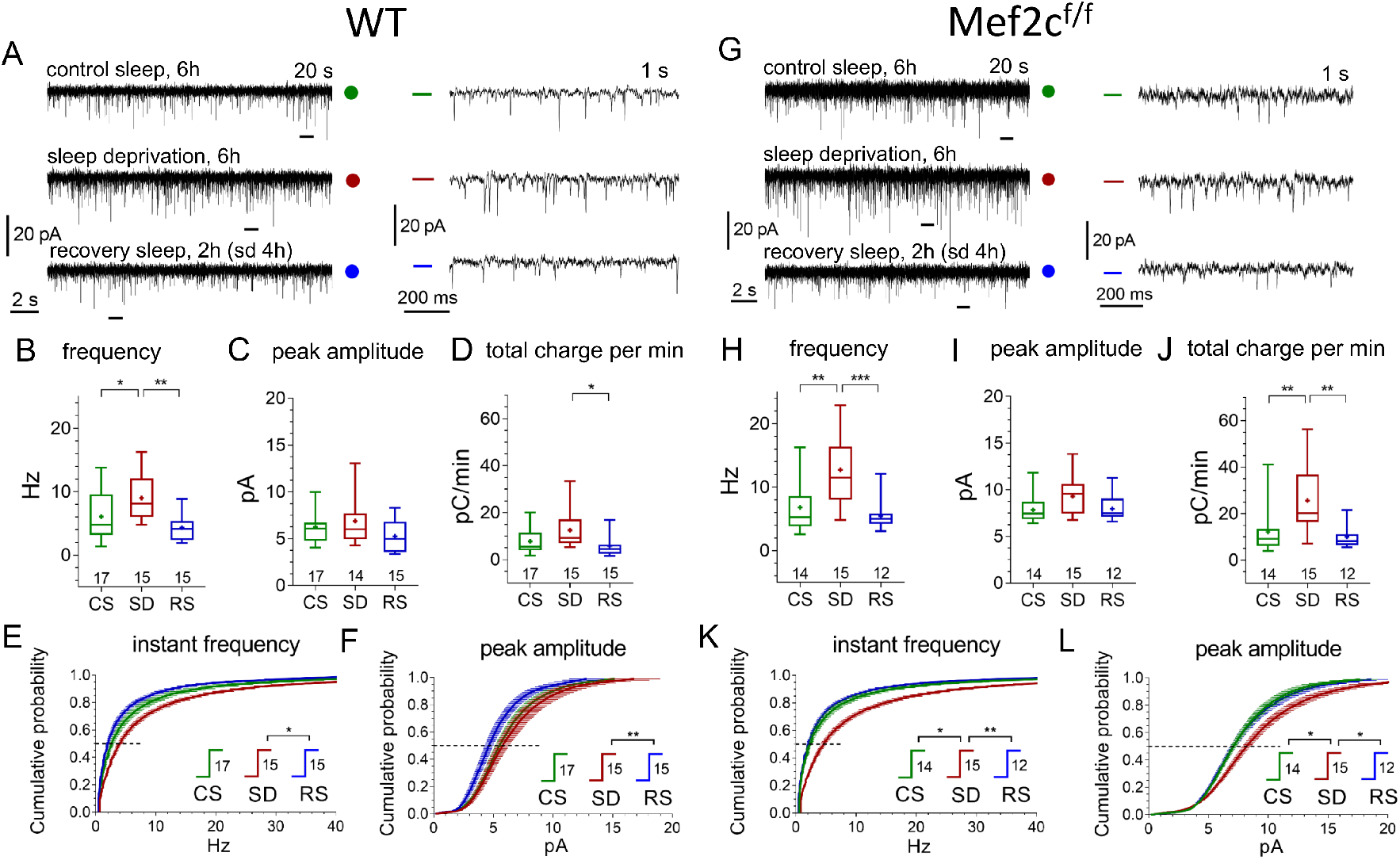
Similar responses of WT and Mef2c^f/f^ mEPSCs to SD and RS conditions. **A, G** Representative recordings of miniature excitatory postsynaptic current (mEPSC) traces (20 s, left panels, and expanded 1s traces on the right corresponding to time bars underneath original traces) obtained for three different experimental sleep/wake conditions, CS (green), SD (red), and RS (blue), for WT mice **(A)** and *Mef2c^f/f^* mice **(G)**. **(B,C,D)** and **(H,I,J)** illustrate mEPSC functional parameters obtained from WT and *Mef2c^f/f^* (number of cells for each condition shown above X axis condition label. Averaged cumulative probability histograms for instant frequency and peak amplitude show increased frequency and amplitude with SD compared to CS or RS for WT **(E, F)** and Mef2c^f/f^ mice **(K, L)**.

**Fig. S6:**
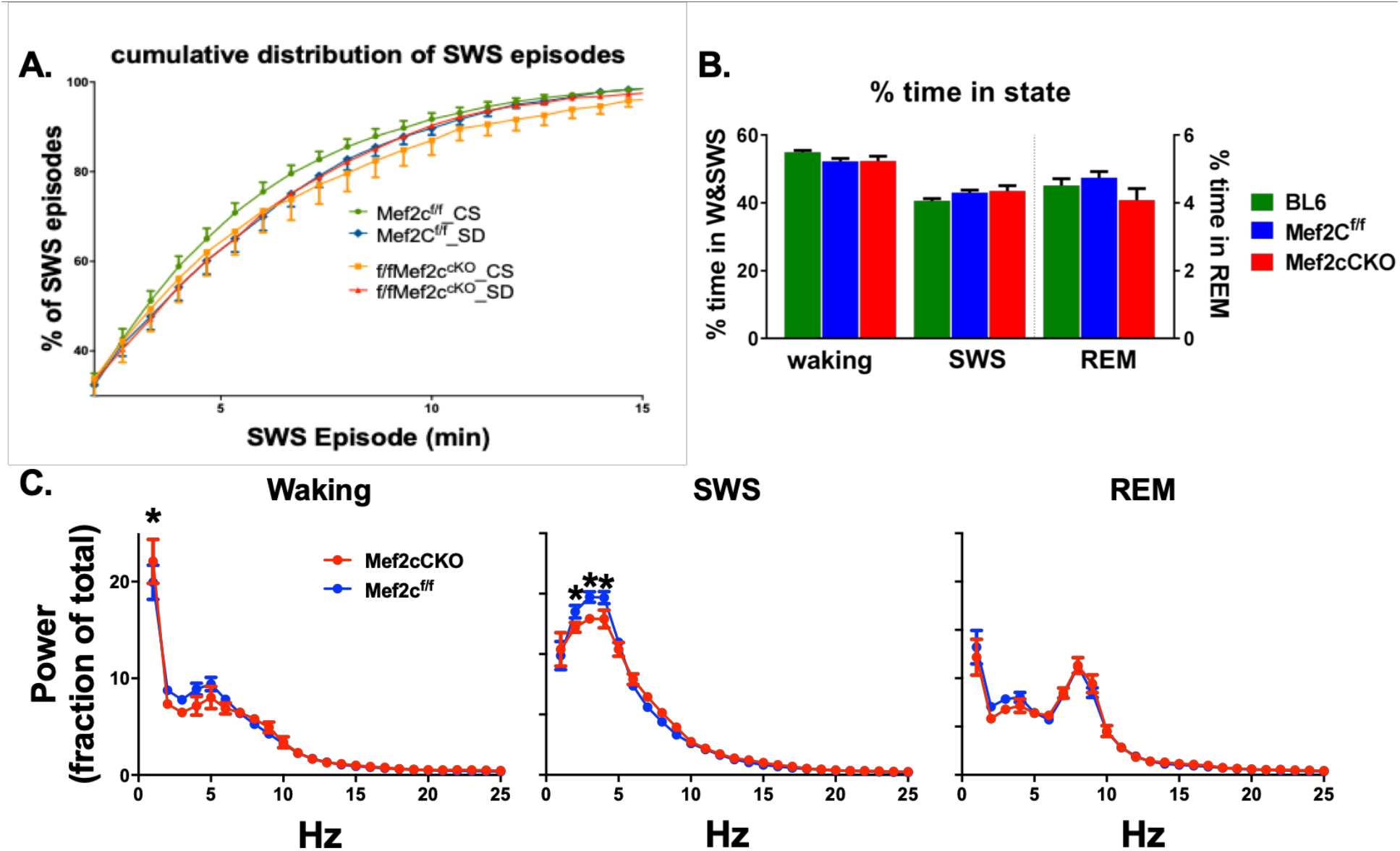
Loss of Mef2c does not influence sleep/waking time, but modestly alters spectral power distribution across states. **(A)** A cumulative histogram of SWS episode durations shows a shift to the right for *Mef2c^f/f^* from CS to SD condition indicative of increased need for sleep. For *Mef2c^cKO^* the SD induced rightward shift is occluded, suggesting high sleep need in CS condition. **(B)** Time in waking, SWS, and REM (right axis) as a percent of total time is unchanged by loss of *Mef2c*. **(C)** Spectral power distribution is modulated by loss of MEF2C function. During waking, relative power in Mef2c^cKO^ mice is shifted to the slower delta range frequencies compared to *Mef2c^f/f^* mice; during SWS, power in the delta range is decreased with loss of MEF2C function; during REM the power distribution is unaffected by genotype. Significant group by frequency differences are indicated by * for *Mef2c^f/f^* and *Mef2c^cKO^* groups.

**Fig. S7.**
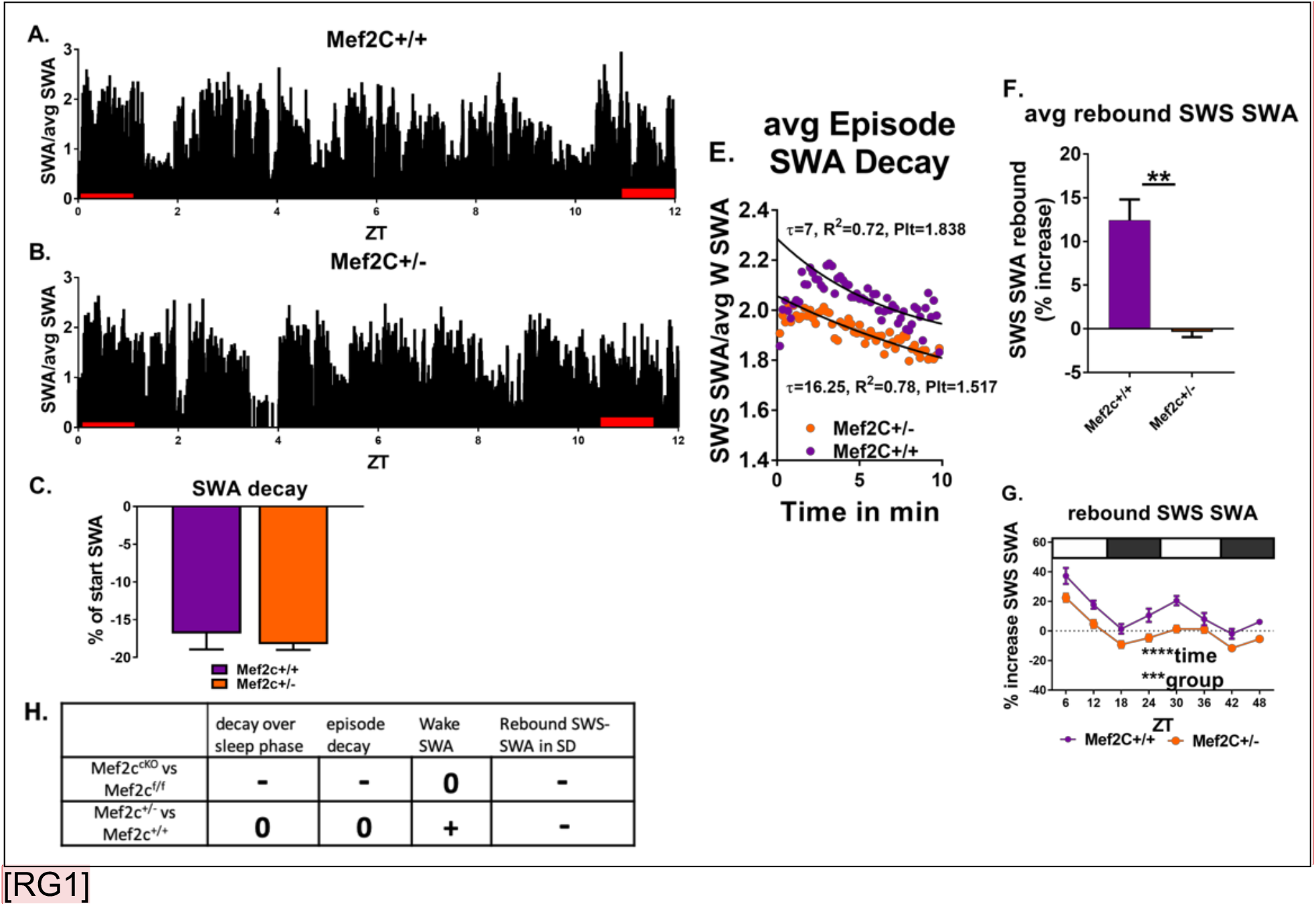
*Mef2c^+/-^* mice resolve sleep need in undisturbed conditions but lack a rebound increase of SWS-SWA power in response to sleep loss. **(A,B)** Average SWA power per 10 second epoch decreases during the sleep phase (ZT=0 to 12h) to the same extent for *Mef2c^+/+^* (n=6) and *Mef2c^+/-^* (n=6) mice (**C**). (**D**) Twenty-four hour average waking SWA power is greater for *Mef2c^+/-^* mice. (**E**) Decay of SWA power (normalized to the average 24h waking SWA) during an average SWS episode is preserved in the *Mef2c^+/-^* mice but with an increased τ, associated with increased sleep need. (**F, G**) Rebound increased SWS-SWA in response to SD is absent in *Mef2c^+/-^* mice. The SD was administered in repeated 6 hour cycles of 4hours SD and 2 hours RS. (**H**) A table showing responses of *Mef2c^cKO^* and *Mef2c^+/-^* compared to their respective controls. (see Fig. 6).

**Fig. S8.**
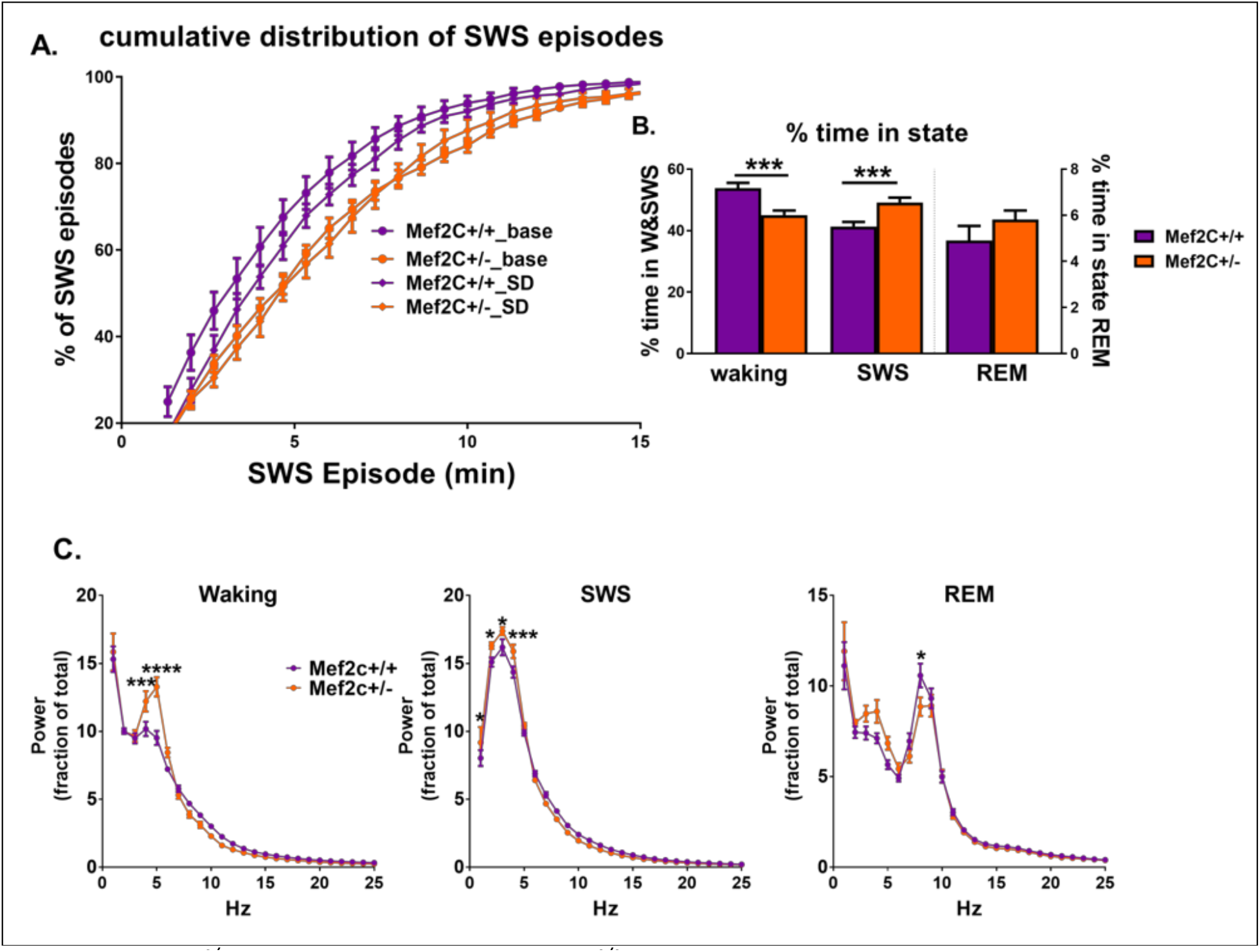
*Mef2c^+/-^* mice are similar to *Mef2c^+/+^* mice with respect to SD evoked increase in SWS episode consolidation. **(A)** A cumulative histogram SWS episode duration showing a generalized shift to the right of the episode duration evoked by SD for both *Mef2c^+/-^* and *Mef2c^+/+^* mice. **(B)** Histograms of relative time spent in waking, SWS or REM show *Mef2c^+/-^* mice have significantly less waking time and increased SWS time under baseline conditions. **(C)** The frequency/power distribution of the EEG shows that *Mef2c^+/-^* mice have relatively more fast delta and slow theta-range power during waking, more delta-range power during SWS, and more mid-theta-range power during REM, than their *Mef2c^+/+^* controls.

**Fig. S9.**
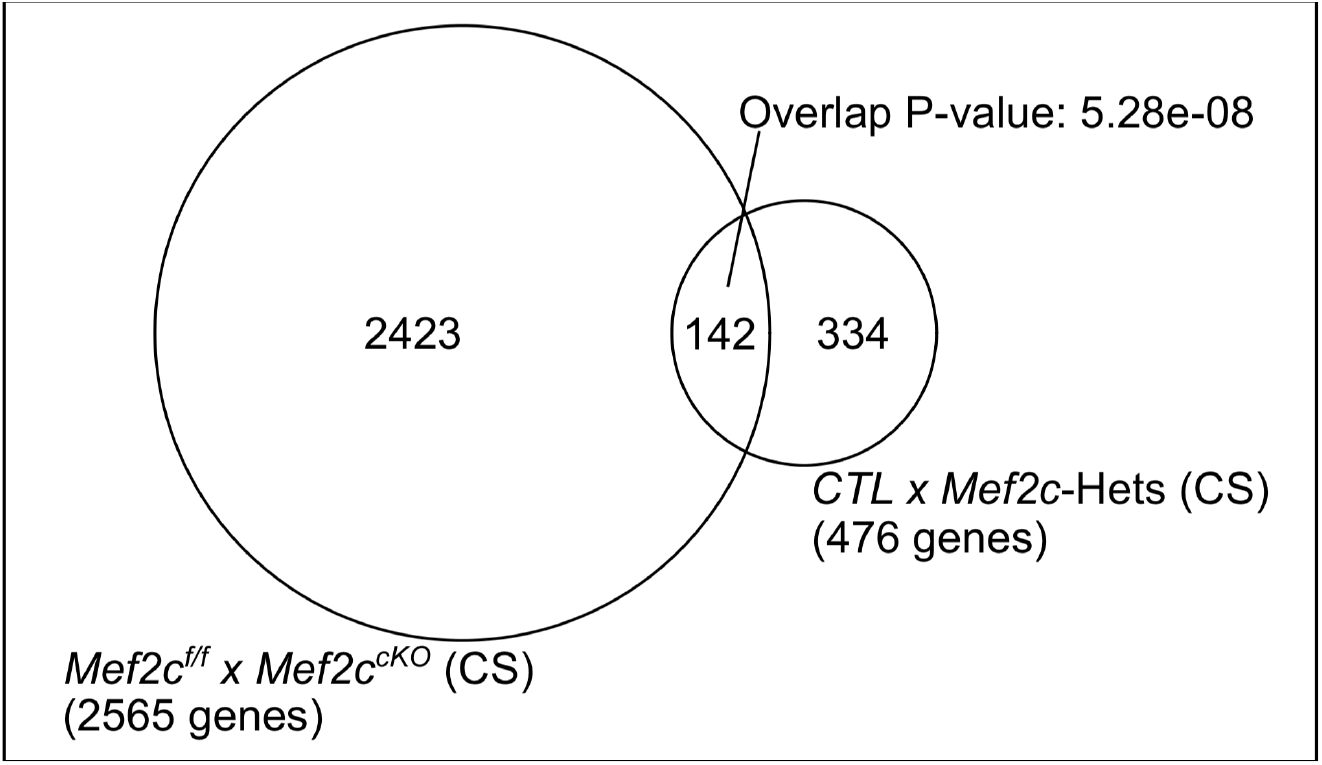
Total of 2565 differentially expressed genes across *Mef2c^f/f^* and *Mef2c^cKO^* under CS conditions. 476 differentially expressed genes from *Mef2c^+/-^* compared to *Mef2c^+/+^* at CS share a significant set of overlapping genes (142; P-value: 5.28e-08, Fisher’s exact test).

**Table S1.**

**Differentially expressed genes (DEG) across sleep states.** (1) DEG for CS to SD, (2) DEG for CS to RS, (3) DEG for SD to RS, (4) list of overlapping and unique genes across sleep states (Fig. 1D) and (5) list of overlapping genes between genes shared by SD and RS against rPRGs (*Tyssowski et al.*) and ERTFs (*Hrvatin et al.*)

**Table S2.**

Overlapping SD genes with glucocorticoid cellular response genes. The GC response genes are a subset (27/44) from MGI GO:0071385 that we identified as being expressed in the frontal cortex.

**Table S3.**

**GO for shared and unique differentially expressed genes (DEG).** (1) GO for genes shared by SD and RS with increased expression against CS, (2) GO for genes shared by SD and RS with decreased expression against CS, (3) GO for genes unique to SD with increased expression against CS, (4) GO for genes unique to SD with decreased expression against CS, (5) GO for genes unique to RS with increased expression against CS and (6) GO for genes unique to RS with decreased expression against CS.

**Table S4.**

**GO for gene modules identified by weighted correlation network analysis (WGCNA)**. (1) Intermodular connectivity across modules, (2) genes associated with each module (3-61) GO for genes within each module.

**Table S5.**

**Differentially expressed genes (DEG) across sleep states for *Mef2c^cKO^*.** (1) DEG for CS to SD for *Mef2c^f/f^* and (2) DEG for CS to SD for *Mef2c^CKO^*.

**Table S6.**

**Functional parameters of mEPSCs** Recordings were obtained *ex Vivo* from anterior cingulate cortex excitatory neurons of Mef2c^f/f^ and Mef2c^cKO^ mice exposed to three different sleep/wake experimental conditions: control sleep 6h (CS), sleep deprivation 6h (SD) and sleep deprivation 4h followed by recovery sleep 2h (RS).

**Table S7.**

**Paired pulse ratio of evoked EPSCs.** P2/P1 was obtained at three different interpulse intervals (20, 50 and 100 ms), obtained in anterior cingulate cortex excitatory neurons of Mef2c^f/f^ and Mef2c^CKO^ mice exposed to three different sleep/wake experimental conditions: control sleep 6h (CS), sleep deprivation 6h (SD) and sleep deprivation 4h followed by recovery sleep 2h (RS).

**Table S8.**

Overlapping genes shared by (a) differentially expressed genes across *Mef2c^f/f^* and *Mef2c^cKO^* at CS and (b) differentially expressed genes across *Mef2c*-Hets and CTL at CS.

## Notes

### Competing Interest Statement

The authors have declared no competing interest.

