## Supplementary material for "An essential role for MEF2C in the cortical response to loss of sleep": Table S6, Table S7

**Supplementary Table 6. Functional parameters of mEPSCs obtained in anterior cingulate cortex excitatory neurons of Mef2c^f/f^ and Mef2c^CKO^ mice exposed to three different sleep/wake experimental conditions: control sleep 6h (CS), sleep deprivation 6h (SD) and sleep deprivation 4h followed by recovery sleep 2h (RS).**

| **mEPSC parameter** | Units | **Mef2c^f/f^** | | | | | | | | | | **Mef2c^CKO^** | | | | | | | | | |
| --- | --- | --- | --- | --- | --- | --- | --- | --- | --- | --- | --- | --- | --- | --- | --- | --- | --- | --- | --- | --- | --- |
|  |  | *Fig.4 B,C, D* | | | | | | | | | | *Fig.4 H,I,J* | | | | | | | | | |
| *From N cell experim. values* |  | N | | Mean ± SEM | ANOVA F, (DFn, DFd) | | ANOVA, Adjusted P value | | | | | N | Mean ± SEM | | ANOVA F, (DFn, DFd) | | ANOVA, Adjusted P value | | | | |
|  |  |  |  |  |  |  | CS | SD | | RS | |  |  |  |  |  | CS | | SD | | RS |
| Frequency, CS | Hz | 14 | | 6.8±1.1 | 11.3  (2,38) | | N/A | 0.002 | | 0.707 | | 14 | 9.0±1.3 | | 0.06  (2,36) | | N/A | | 0.96 | | 0.996 |
| Frequency, SD | Hz | 15 | | 12.7±1.4 |  |  | 0.002 | N/A | | 0.0003 | | 12 | 8.5±1.4 | |  |  | 0.96 | | N/A | | 0.94 |
| Frequency, RS | Hz | 12 | | 5.4±0.7 |  |  | 0.707 | 0.0003 | | N/A | | 13 | 9.2±1.6 | |  |  | 0.996 | | 0.94 | | N/A |
| Peak ampl., CS | pA | 14 | | 7.9±0.4 | 3.44  (2,38) | | N/A | 0.058 | | 0.98 | | 14 | 9.0±0.5 | | 0.43  (2,36) | | N/A | | 0.99 | | 0.66 |
| Peak ampl., SD | pA | 15 | | 9.3±0.5 |  |  | 0.058 | N/A | | 0.106 | | 12 | 9.1±0.6 | |  |  | 0.99 | | N/A | | 0.76 |
| Peak ampl., RS | pA | 12 | | 8.0±0.4 |  |  | 0.98 | 0.106 | | N/A | | 13 | 9.8±0.9 | |  |  | 0.66 | | 0.76 | | N/A |
| Charge/min, CS | pC/min | 14 | | 12.1±2.6 | 9.05  (2,38) | | N/A | 0.0041 | | 0.86 | | 14 | 16.6±3.2 | | 0.32  (2,36) | | N/A | | 0.96 | | 0.86 |
| Charge/min, SD | pC/min | 15 | | 25.6±3.6 |  |  | 0.0041 | N/A | | 0.0014 | | 12 | 15.3±1.4 | |  |  | 0.96 | | N/A | | 0.71 |
| Charge/min, RS | pC/min | 12 | | 9.9±1.5 |  |  | 0.86 | 0.0014 | | N/A | | 13 | 19.2±4.1 | |  |  | 0.86 | | 0.71 | | N/A |
|  |  | *Fig.4 B,C, D* | | | | | | | | | | *Fig.4 H,I,J* | | | | | | | | | |
| *From N cell experim. values* |  | N | Median | | | 25% -tile | 75% -tile | | Min value | | Max value | N | | Median | | 25% -tile | 75% -tile | Min value | | Max value | |
| Frequency, CS | Hz | 14 | 5.29 | | | 3.83 | 8.57 | | 2.58 | | 16.3 | 14 | | 8.12 | | 5.22 | 11.8 | 1.16 | | 17.5 | |
| Frequency, SD | Hz | 15 | 11.5 | | | 8.05 | 16.4 | | 4.80 | | 22.9 | 12 | | 6.12 | | 4.91 | 12.8 | 2.79 | | 19.0 | |
| Frequency, RS | Hz | 12 | 5.03 | | | 4.20 | 5.82 | | 3.05 | | 12.1 | 13 | | 8.68 | | 4.06 | 12.5 | 2.70 | | 21.9 | |
| Peak ampl., CS | pA | 14 | 7.46 | | | 6.85 | 8.73 | | 6.43 | | 11.8 | 14 | | 8.81 | | 7.46 | 9.84 | 6.40 | | 13.1 | |
| Peak ampl., SD | pA | 15 | 9.56 | | | 7.44 | 10.6 | | 6.75 | | 13.8 | 12 | | 8.59 | | 7.87 | 9.92 | 6.93 | | 14.5 | |
| Peak ampl., RS | pA | 12 | 7.50 | | | 7.11 | 9.07 | | 6.57 | | 11.3 | 13 | | 8.43 | | 7.76 | 12.5 | 5.16 | | 17.0 | |
| Charge/min, CS | pC/min | 14 | 9.17 | | | 6.17 | 13.4 | | 3.92 | | 41.1 | 14 | | 13.7 | | 9.71 | 19.7 | 1.95 | | 49.9 | |
| Charge/min, SD | pC/min | 15 | 20.2 | | | 16.4 | 36.8 | | 7.14 | | 56.3 | 12 | | 11.5 | | 8.38 | 19.5 | 4.92 | | 40.7 | |
| Charge/min, RS | pC/min | 12 | 8.22 | | | 6.42 | 11.1 | | 5.51 | | 21.6 | 13 | | 15.2 | | 9.71 | 28.4 | 3.60 | | 51.5 | |
|  |  | *Fig.4 E,F* | | | | | | | | | | *Fig.4 K,L* | | | | | | | | | |
| *From N averaged cumul. histograms* |  | N | | Average median  ± SEM | ANOVA F, (DFn, DFd) | | ANOVA, Adjusted P value | | | | | N | Average median  ± SEM | | ANOVA F, (DFn, DFd) | | ANOVA, Adjusted P value | | | | |
|  |  |  |  |  |  |  | CS | SD | | RS | |  |  |  |  |  | CS | | SD | | RS |
| Inst. freq., CS | Hz | 14 | | 2.4±0.4 | 6.18  (2,297) | | N/A | 0.020 | | 0.82 | | 14 | 3.0 ±0.4 | | 0.07  (2,294) | | N/A | | 0.99 | | 0.96 |
| Inst. freq., SD | Hz | 15 | | 4.6±0.5 |  |  | 0.020 | N/A | | 0.0031 | | 12 | 3.0±0.5 | |  |  | 0.99 | | N/A | | 0.93 |
| Inst. freq., RS | Hz | 12 | | 2.1±0.3 |  |  | 0.82 | 0.0031 | | N/A | | 13 | 3.4±0.6 | |  |  | 0.96 | | 0.93 | | N/A |
| Peak ampl., CS | pA | 14 | | 7.1±0.3 | 4.37  (2,297) | | N/A | 0.022 | | 0.97 | | 14 | 7.8±0.3 | | 0.95  (2,297) | | N/A | | 0.96 | | 0.39 |
| Peak ampl., SD | pA | 15 | | 8.2±0.4 |  |  | 0.022 | N/A | | 0.040 | | 12 | 7.9±0.3 | |  |  | 0.96 | | N/A | | 0.55 |
| Peak ampl., RS | pA | 12 | | 7.1±0.3 |  |  | 0.97 | 0.040 | | N/A | | 13 | 8.1±0.6 | |  |  | 0.39 | | 0.55 | | N/A |

**Supplementary Table 7. Paired pulse ratio of evoked EPSCs at three different interpulse intervals (20, 50 and 100 ms), obtained in anterior cingulate cortex excitatory neurons of Mef2c^f/f^ and Mef2c^CKO^ mice exposed to three different sleep/wake experimental conditions: control sleep 6h (CS), sleep deprivation 6h (SD) and sleep deprivation 4h followed by recovery sleep 2h (RS).**

| **P2/P1 ratio** | | **Mef2c^f/f^** | | | | | | | | **Mef2c^CKO^** | | | | | | | | |
| --- | --- | --- | --- | --- | --- | --- | --- | --- | --- | --- | --- | --- | --- | --- | --- | --- | --- | --- |
|  | | *Fig.5 B* | | | | | | | | *Fig.5 D* | | | | | | | | |
| *From N cell experimental values* | | N | Mean ± SEM | ANOVA F, (DFn, DFd) | ANOVA, Adjusted P value | | | | | N | Mean ± SEM | ANOVA F, (DFn, DFd) | | ANOVA, Adjusted P value | | | | |
|  |  |  |  |  | CS | SD | | RS | |  |  |  |  | CS | SD | | RS | |
| 20 ms interpuls time, CS | | 8 | 0.72±0.07 | 3.27  (2,20) | N/A | 0.42 | | 0.047 | | 9 | 0.88±0.09 | 1.54  (2,20) | | N/A | 0.22 | | 0.88 | |
| 20 ms interpuls time, SD | | 7 | 0.87±0.07 |  | 0.42 | N/A | | 0.48 | | 9 | 1.04±0.04 |  |  | 0.22 | N/A | | 0.60 | |
| 20 ms interpuls time, RS | | 8 | 1.00±0.09 |  | 0.047 | 0.48 | | N/A | | 5 | 0.93±0.06 |  |  | 0.88 | 0.60 | | N/A | |
| 50 ms interpuls time, CS | | 8 | 0.77±0.04 | 4.71  (2,21) | N/A | 0.16 | | 0.017 | | 9 | 0.86±0.06 | 1.42  (2,21) | | N/A | 0.24 | | 0.65 | |
| 50 ms interpuls time, SD | | 8 | 0.93±0.07 |  | 0.16 | N/A | | 0.50 | | 9 | 1.02±0.07 |  |  | 0.24 | N/A | | 0.82 | |
| 50 ms interpuls time, RS | | 8 | 1.03±0.07 |  | 0.017 | 0.50 | | N/A | | 6 | 0.96±0.09 |  |  | 0.65 | 0.82 | | N/A | |
| 100 ms interpuls time, CS | | 10 | 0.75±0.04 | 1.86  (2,23) | N/A | 0.21 | | 0.31 | | 9 | 0.89±0.06 | 2.32  (2,22) | | N/A | 0.11 | | 0.34 | |
| 100 ms interpuls time, SD | | 8 | 0.89±0.06 |  | 0.21 | N/A | | 0.97 | | 9 | 1.06±0.05 |  |  | 0.11 | N/A | | 0.86 | |
| 100 ms interpuls time, RS | | 8 | 0.87±0.06 |  | 0.31 | 0.0.97 | | N/A | | 7 | 1.01±0.06 |  |  | 0.34 | 0.86 | | N/A | |
|  |  | *Fig.5 B* | | | | | | | | *Fig.5 D* | | | | | | | | |
| *From N cell experimental values* | | N | Median | 25% -tile | 75% -tile | | Min value | | Max value | N | Median | | 25% -tile | 75% -tile | | Min value | | Max value |
| 20 ms interpuls time, CS | | 8 | 0.71 | 0.62 | 0.86 | | 0.38 | | 1.06 | 9 | 0.95 | | 0.61 | 1.09 | | 0.46 | | 1.22 |
| 20 ms interpuls time, SD | | 7 | 0.93 | 0.73 | 0.99 | | 0.52 | | 1.02 | 9 | 1.06 | | 0.94 | 1.16 | | 0.83 | | 1.22 |
| 20 ms interpuls time, RS | | 8 | 1.06 | 0.77 | 1.19 | | 0.57 | | 1.31 | 5 | 0.89 | | 0.82 | 1.07 | | 0.79 | | 1.14 |
| 50 ms interpuls time, CS | | 8 | 0.81 | 0.66 | 0.86 | | 0.54 | | 0.89 | 9 | 0.89 | | 0.71 | 0.94 | | 0.66 | | 1.20 |
| 50 ms interpuls time, SD | | 8 | 0.96 | 0.75 | 1.11 | | 0.63 | | 1.14 | 9 | 1.00 | | 0.91 | 1.11 | | 0.71 | | 1.44 |
| 50 ms interpuls time, RS | | 8 | 1.01 | 0.89 | 1.17 | | 0.77 | | 1.34 | 6 | 0.98 | | 0.75 | 1.09 | | 0.70 | | 1.28 |
| 100 ms interpuls time, CS | | 10 | 0.72 | 0.65 | 0.89 | | 0.55 | | 0.97 | 9 | 0.91 | | 0.74 | 0.98 | | 0.67 | | 1.24 |
| 100 ms interpuls time, SD | | 8 | 0.92 | 0.73 | 1.06 | | 0.59 | | 1.07 | 9 | 1.06 | | 0.91 | 1.18 | | 0.85 | | 1.29 |
| 100 ms interpuls time, RS | | 8 | 0.89 | 0.73 | 0.98 | | 0.60 | | 1.18 | 7 | 1.01 | | 0.94 | 1.13 | | 0.75 | | 1.26 |
